# Dynamic centriolar relocalization of Polo kinase and Centrobin in early mitosis primes centrosome asymmetry in fly neural stem cells

**DOI:** 10.1101/249375

**Authors:** Emmanuel Gallaud, Anjana Ramdas Nair, Nicole Horsley, Arnaud Monnard, Priyanka Singh, Tri Thanh Pham, David Salvador Garcia, Alexia Ferrand, Clemens Cabernard

## Abstract

Centrosomes, the main microtubule organizing centers (MTOCs) of metazoan cells, contain an older ‘mother’ and a younger ‘daughter’ centriole. Stem cells either inherit the mother or daughter centriole-containing centrosome, providing a possible mechanism for biased delivery of cell fate determinants. However, the dynamics and mechanisms regulating centrosome asymmetry and biased centrosome segregation are unclear. Using 3D-Structured Illumination Microscopy (3D-SIM) and live cell imaging we show that in fly neural stem cells (neuroblasts) the mitotic kinase Polo and its centriolar protein substrate Centrobin (Cnb) dynamically relocalize from the mother to the daughter centriole during mitosis. This mechanism generates a centrosome, containing two molecularly distinct centrioles by telophase. Cnb’s timely relocalization is regulated by Polo-mediated phosphorylation whereas Polo’s daughter centriole enrichment requires both Wdr62 and Cnb. Based on optogenetic protein mislocalization experiments we propose that the establishment of centriole asymmetry in mitosis primes biased interphase MTOC activity, necessary for correct spindle orientation.

## Introduction

Centrosomes consist of a pair of centrioles, embedded in structured layers of pericentriolar material (PCM) ^1^. During interphase of each cell cycle a single ‘daughter’ centriole is formed around a central cartwheel at a right angle to the existing older ‘mother’ centriole ^2-4^. Based on this replication cycle, centrioles - and thereby centrosomes – have an intrinsic age asymmetry. Centrosome asymmetry is also manifested in the unequal clustering of proteins or mRNA ^5-7^. Many metazoan cells recognize centrosomal asymmetry as a cue for biased centrosome segregation, providing a possible mechanism to determine, or influence, cell fate decisions ^8^. For instance, vertebrate neural stem cells and *Drosophila* male germline stem cells both retain the mother centriole-containing centrosome (mother centrosome hereafter) ^9,10^, while *Drosophila* female germline or neural stem cells, called neuroblasts, inherit the daughter centriole-containing centrosome (daughter centrosome hereafter) ^11-13^.

In *Drosophila* male germline or neural stem cells, asymmetric centrosome function mediates spindle orientation ^10,14^. Correct spindle orientation is necessary for stem cell cycle progression, stem cell homeostasis and differentiation ^15,16^. However, the mechanisms establishing functional centrosome asymmetry are incompletely understood. Furthermore, how centrosome asymmetry affects biased centrosome segregation remains elusive.

Here, we use *Drosophila* neuroblasts to investigate the spatiotemporal mechanisms underlying the establishment of centrosome asymmetry *in vivo*. Neuroblast centrosomes are highly asymmetric in interphase: one centrosome forms an active MTOC, while its sibling remains inactive until entry into mitosis ^12,14,17^. The active interphase MTOC contains the daughter centriole, identifiable with the orthologue of the human daughter centriole-specific protein Cnb (Cnb^+^) ^11^. This biased MTOC activity is regulated by the mitotic kinase Polo (Plk1 in vertebrates). Polo phosphorylates Cnb, necessary to maintain an active MTOC, tethering the daughter centriole-containing centrosome to the apical interphase cortex (the apical centrosome hereafter) ^18^. Apical centrosome tethering predetermines the alignment of the mitotic spindle along the intrinsic apical-basal polarity axis. Furthermore, this cortical association ensures that the daughter centrosome is inherited by the self-renewing neuroblast ^11,18^. Polo localization on the apical centrosome is maintained by the microcephaly associated protein Wdr62 ^19^. The mother centrosome, separating from the daughter centrosome in interphase, downregulates Polo and MTOC activity through Pericentrin (PCNT)-like protein (Plp) and Bld10 (Cep135 in vertebrates) ^20,21^. The lack of MTOC activity prevents the mother centrosome from engaging with the apical cell cortex; it randomly migrates through the cytoplasm until centrosome maturation in prophase establishes a second MTOC near the basal cortex (called the basal centrosome hereafter), ensuring its segregation into the differentiating ganglion mother cell (GMC). Later in mitosis, the mother centrosome also accumulates Cnb ^12,14,17,20^ (and Supplementary Fig.1a).

Although several centrosomal proteins have been described to be enriched on either the mother or daughter centrosome in *Drosophila* interphase neuroblasts ^11,19,22^ or human cells ^5^, it is unknown when and how centrosomes acquire their unique molecular identity to determine biased MTOC activity, and thus correct spindle orientation. Here, we show that centrosome asymmetry is primed in early mitosis by dynamically relocalizing Polo and Cnb from the older mother to the younger daughter centriole, while selectively retaining Plp on the mother centriole. We further show that priming centrosome asymmetry in mitosis is necessary to establish molecularly distinct centrosomes, asymmetric MTOC activity and centrosome positioning.

## Results

### Neuroblast centriole duplication starts in interphase and completes in mitosis

To determine the onset of centrosome asymmetry establishment in larval neuroblast, we first investigated the centriole replication cycle (Supplementary Fig.1c). In vertebrate cells, centrioles replicate in interphase and convert to functional centrosomes during mitosis (reviewed in ^3,4,23^) but it is unclear whether this also applies to fly neuroblast. We used 3D-Structured Illumination Microscopy (3D-SIM), which has approximately twice the spatial resolution of standard confocal microscopy, and stained third instar neuroblasts with known centriolar and centrosomal markers. For all the 3D-SIM experiments, the cell cycle stages were determined based on the organization of the microtubule network (Supplementary Fig.1b). We used Asl in conjunction with Sas-6 to determine the onset of cartwheel duplication and centriole conversion during the neuroblast cell cycle (Supplementary Fig.1c). Consistent with previous reports ^1,24-26^ we found that Sas-6 was localized to the centriolar cartwheel whereas Asterless (Asl) surrounded the centriolar wall (Supplementary Fig.1d). Asl has been shown to extend from the core centriolar region into the adjacent PCM and sequentially loads onto the new centriole during centriole-to-centrosome conversion (also referred to as mitotic centriole conversion), a mechanism generating a centriole-duplication and PCM-assembly competent centrosome ^25,27,28^. Apical and basal interphase neuroblast centrosomes contained two Sas-6^+^ cartwheels but only one Asl^+^ centriole (Supplementary Fig.1d, yellow arrowhead). From prometaphase onwards, Asl gradually appeared around the second cartwheel to form a pair of fully formed centrioles. In telophase, centrioles seemed to lose their orthogonal conformation, possibly due to disengagement before migration. Cartwheels started to duplicate in late telophase, manifested in the appearance of a third Sas-6 positive cartwheel (blue arrowhead in Supplementary Fig.1d). Based on these data we conclude that in third instar larval neuroblasts centriolar cartwheels are duplicated in early interphase, forming a new procentriole. This procentriole subsequently converts into a mature centriole during mitosis through progressive loading of Asl. Thus, by the end of telophase, both neuroblast centrosomes contain an older mother and younger daughter centriole, separating in the following early interphase.

### Asymmetric Cnb localization is established in early mitosis through dynamic exclusion from the mother centriole and enrichment on the daughter centriole

Molecular and functional centrosome asymmetry is detectable in interphase neuroblasts but when and how this asymmetry is established is unclear (Supplementary Fig.1c). To this end, we analyzed the localization of YFP::Cnb ^11^ with 3D-SIM throughout mitosis. As expected, YFP::Cnb was localized with Asl on the active, apical centrosome in interphase neuroblasts but absent on the basal interphase centrosome (Fig. 1a-d). To our surprise, we also found apical - but never basal - prophase and prometaphase centrosomes where Cnb was localized on both centrioles (green arrowheads and bars in Fig. 1b & Fig. 1g). However, Cnb was predominantly localized on one centriole only from metaphase onward (brown arrowheads and bars in Fig. 1b & Fig. 1g). On the basal centrosome, Cnb appeared in prophase and was consistently localized to a single centriole in all subsequent mitotic stages (Fig. 1d, g). Since Asl sequentially loads onto the forming daughter centriole ^25,29^, we tested whether Asl can be used as an independent marker for centriolar age. To this end, we calculated the Asl intensity ratio between both centrioles (see methods) – on the apical and basal centrosome - for all mitotic stages where we could find a clear Cnb asymmetry (Asl intensity ratio of Cnb^+^/Cnb^−^ from prometaphase until telophase). These calculations revealed a clear Asl intensity asymmetry with the Cnb^+^ centriole always containing less Asl and the Cnb^−^ more Asl (Fig. 1e).

**Fig. 1:**
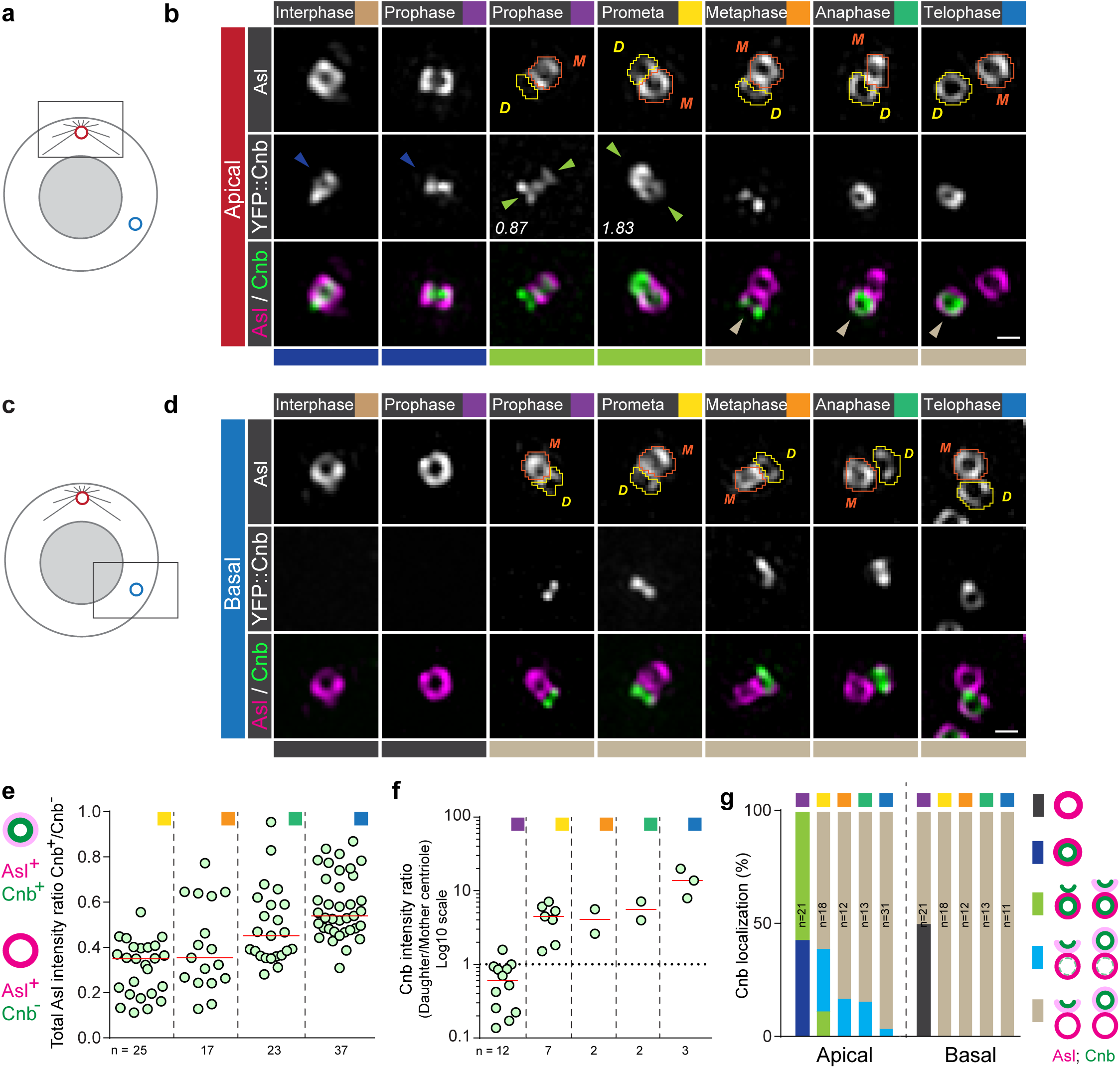
Cnb relocalizes from the mother to the daughter centriole in early mitosis on the apical centrosome. How centriole duplication and molecular asymmetry are coupled is unclear for both the apical **(a)** and basal **(c)** centrosome. Representative 3D-SIM images of apical **(b)** and basal **(d)** third instar neuroblast centrosomes, expressing YFP::Cnb (middle row; white. Green in merge) and stained for Asl (Top row; white. Magenta in merge). Orange and yellow shapes highlight mother and daughter centrioles, respectively and were used to measure signal intensities. The numbers indicate the total Cnb asymmetry ratios (Daughter/Mother centriole). Colored arrowheads and bars underneath the images highlight the different stages shown in (g). **(e)** For prometaphase to telophase centrosomes (apical and basal combined), containing a single Cnb^+^ centriole, total Asl intensity of the Cnb^+^ (presumably the daughter) centriole was divided by the total Asl intensity of the Cnb^−^ (presumably the mother) centriole. Medians are shown with a red horizontal line. **(f)** Scatter plot showing total Cnb intensity of the daughter centriole (less Asl), divided by total Cnb intensity on the mother centriole (more Asl). Only apical centrioles containing Cnb on both centrioles were measured. **(g)** Graph showing the timeline of Cnb’s localization dynamics on the apical and basal centrosome: the bars show the percentage of neuroblasts containing an apical centrosome containing one centriole Cnb^+^ (dark blue), a basal centrosome containing one centriole without Cnb (dark grey), a centrosome with Cnb on both centrioles (transition stage with a Daughter/Mother ratio < 2; light green), predominant Cnb localization on the daughter centriole (strong asymmetry with a Daughter/Mother ratio between 2 and 10; light blue) or in which Cnb is only present on the daughter centriole (complete asymmetry with a Daughter/Mother ratio > 10; light brown) at defined mitotic stages. For this and all subsequent cartoons: closed and open circles represent established mother and forming daughter centrioles, respectively. Cell cycle stages are indicated with colored boxes. Scale bar is 0.3 μm. The data presented here were obtained from five independent experiments.

Using the Asl intensity ratio as a method to distinguish between mother and daughter centrioles, we next correlated Cnb localization with centriolar age at all mitotic stages. We found that in prophase – when Cnb was detectable on both centrioles – Cnb was predominantly associated with the centriole containing more Asl (the mother centriole). However, during prometaphase, more Cnb was localized on the centriole containing less Asl (the daughter centriole). Cnb was sometimes visible before Asl was robustly recruited to the daughter centriole (green arrowheads in second column of Fig. 1b). From metaphase until mitosis exit, Cnb was strongly enriched or exclusively present on the daughter centriole (brown bars and arrowheads Fig. 1b, d, f, g).

From these data, we conclude that neuroblast centrosomes generate two molecularly distinct centrioles during early mitosis. The dynamics generating this centriole asymmetry differ between the apical and basal centrosomes: on the apical centrosome, Cnb is initially only present on the mother centriole before appearing on the daughter, and disappearing on the mother centriole. In contrast, Cnb directly appears on the daughter centriole of the basal centrosome. This establishment of molecular centriole asymmetry occurs during the centriole-to-centrosome conversion period.

### The daughter centriole’s Cnb partially originates from the mother centriole

We next investigated Cnb relocalization dynamics, considering the following two non-exclusive hypotheses: (1) Cnb could directly relocalize from the mother to the newly forming daughter centriole during mitosis. (2) Alternatively, Cnb could be downregulated on the mother and upregulated on the daughter centriole during mitosis, implying that newly recruited Cnb contributes to the apparent relocalization pattern (Fig. 2a). To distinguish between these scenarios, we needed to determine the origin of the daughter centriole specific Cnb pool. To this end, we first performed live cell imaging of endogenously tagged Cnb::EGFP (see methods) in conjunction with the mitotic spindle marker mCherry::Jupiter ^16^. We found that in late interphase, prior to mitotic entry, Cnb was strongly localized on the apical neuroblast centrosome. At this cell cycle stage, the apical centrosome only consists of a single Asl^+^ mother centriole (Supplementary Fig.1d). Subsequently, Cnb got downregulated as the neuroblast entered mitosis and Cnb levels were lowest between prometaphase and anaphase. Cnb intensity then increased again from anaphase onward (Fig. 2b, c). To test whether daughter centriole Cnb originates from the mother centriole, or is recruited from other sources, we performed Fluorescence Recovery After Photobleaching (FRAP) experiments. Bleaching Cnb on the apical centrosome in late interphase or early prophase extinguished Cnb fluorescence, which only recovered from anaphase onward (Fig. 2d-f). We also tagged Cnb endogenously with mDendra2 (see also below) but the signal was too low to perform photoconversion experiments. Regardless, the lack of Cnb fluorescence recovery during mitosis indicates that very little to no new Cnb is recruited to the apical centrosome prior to anaphase. Recovery of Cnb after anaphase onset suggests the existence of a Cnb protein pool different from the Cnb initially localized to the apical mother centriole. Taken together, we conclude that Cnb on the daughter centriole is composed of Cnb originating from the mother centriole in early mitosis and newly recruited Cnb from anaphase onward (Fig. 2g).

**Fig. 2:**
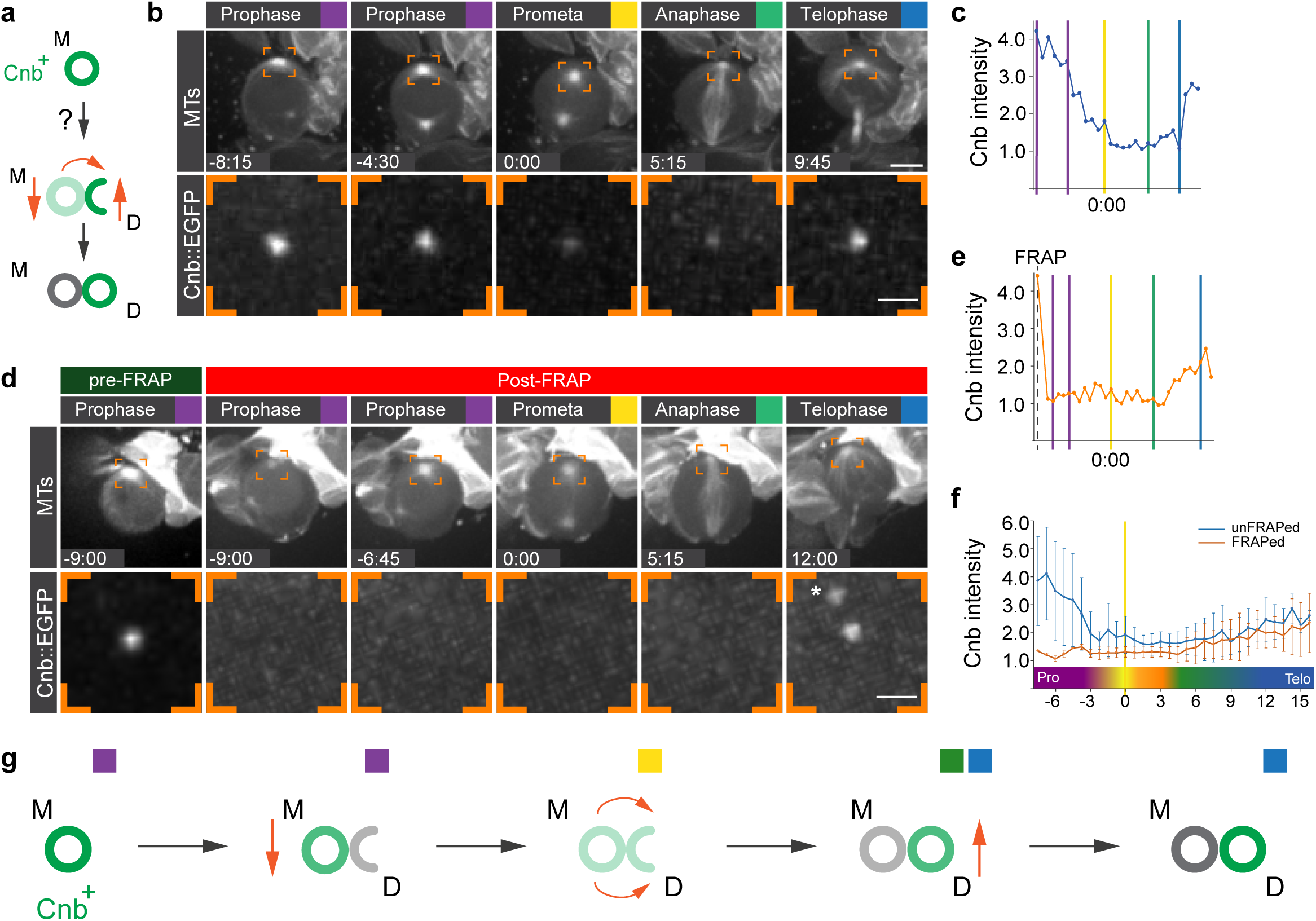
Cnb localized on the daughter centriole partially originates from the mother centriole. **(a)** Dynamic mother-daughter centriole relocalization of Cnb could be either due to a direct transfer mechanism (orange curved arrow) or through up- and downregulation (vertical orange arrows). Representative unFRAPed **(b)** and FRAPed **(d)** wild type neuroblast expressing endogenously tagged Cnb::EGFP (white; bottom row) together with the microtubule (MT) marker mCherry::Jupiter (white; top row). The orange brackets highlight the apical centrosome where Cnb::EGFP (bottom row) is measured. The asterisk refers to an unspecific Cnb::EGFP aggregate. Intensity profile of the unFRAPed **(c)** and FRAPped **(e)** apical Cnb::EGFP signal of the neuroblasts shown in (b) and (d). Colored vertical bars indicate specific cell cycle stages. The vertical dashed line refers to the timepoint when bleaching was performed. Cnb fluorescence was normalized against cytoplasmic EGFP levels. **(f)** Mean intensity plot of 10 unFRAPed and frapped apical centrosomes. Error bars indicate standard deviation of the mean. **(g)** Graphical summary for apical Cnb: Cnb levels decrease during prometaphase. The remaining apical Cnb transfers from the mother to the daughter centriole until anaphase. From anaphase onward, Cnb levels increase again through recruitment of new Cnb. Time scale is mm:ss. Scale bar in (b) and (d) is 5μm (top row) and 1μm (bottom row). The data presented here were obtained from three independent experiments.

### Polo dependent phosphorylation of Cnb is necessary for a timely relocalization of Cnb from the mother to the daughter centriole

Previously, it was shown that Cnb is a substrate of Polo ^18^. We thus tested whether Cnb’s dynamic relocalization depends on Polo phosphorylation by analyzing YFP::Cnb localization in hypomorphic *polo* mutant neuroblasts (*polo^16-1^*/*polo^1^*). In addition, we analyzed the localization of YFP::Cnb^T4A,T9A,S82A^, a mutant version of Cnb in which all three consensus phosphorylation sites for Polo were substituted by alanine ^18^, in *cnb* mutant neuroblasts. Since we cannot accurately distinguish between apical and basal centrosomes in *polo* mutants, or *cnb* mutants expressing YFP::Cnb^T4A,T9A,S82A^, we will refer to them as centrosome 1 and centrosome 2. In contrast to prophase wild type or control (*polo/+* heterozygotes) neuroblasts, showing no Cnb on the mother centriole of the basal centrosome (Fig. 1; Supplementary Fig. 2c), we found *polo* mutant neuroblasts containing weak Cnb on the mother centriole of both prophase centrosomes (44%, light blue and green arrowheads and bars, centrosome 2, Supplementary Fig.2a, b, d). In prometaphase and metaphase neuroblasts Cnb appeared on both centrioles on centrosome 1 and 2 (prometaphase: 14.8%; n = 27; metaphase: 18.8%; n =16) (light blue and green arrowheads and bars, centrosome 2, Supplementary Fig.2a, b and D) but from anaphase onward was predominantly localized on the mother centriole. Taken together, Cnb relocalization occurs but is delayed in *polo* hypomorphic mutant neuroblasts (Supplementary Fig.2a-d).

A similar, albeit stronger phenotype was observed in *cnb* mutant neuroblasts expressing YFP::Cnb^T4A,T9A,S82A^; neuroblasts containing Cnb^+^ mother centrioles on both centrosomes were found for all mitotic stages. Similar to Cnb in *polo* mutant neuroblasts, phosphomutant Cnb was detectable on both centrosomes in early prophase neuroblasts (e.g centrosome 2: 68.6%; n = 19; light blue and green arrowheads and bars) (Fig. 3a-c). Due to their resemblance to apical wild type centrosomes in regard of Cnb localization, we refer to these centrosomes as “apical-like”. In most wild type neuroblasts, Cnb was relocalized by metaphase but in *cnb* mutant neuroblasts expressing YFP::Cnb^T4A,T9A,S82A^, 71.4% (n = 7; light blue, centrosome 1) of analyzed neuroblasts show incomplete Cnb relocalization on one centrosome by telophase (Fig. 3a-c).

**Fig. 3:**
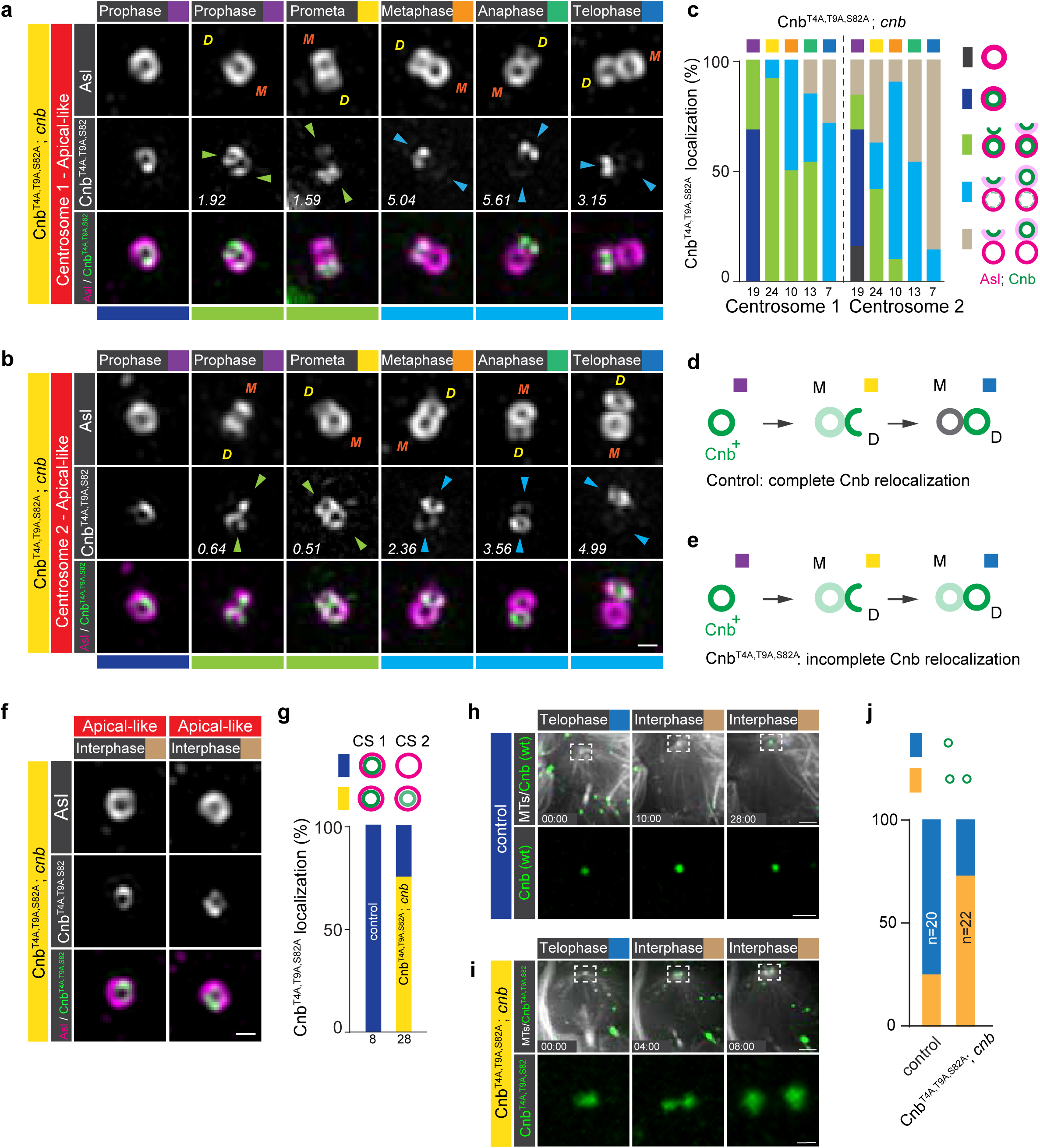
Cnb’s relocalization from the mother to the daughter centriole is controlled by Polo-dependent phosphorylation. Expression of YFP::Cnb^T4A,T9A,S82A^ in *cnb* mutant neuroblasts generates two “apical-like” (in respect of Cnb localization) centrosomes. Because we cannot distinguish between “apical” and “basal” centrosome anymore, they are labelled centrosome 1 and 2, respectively. Representative 3D-SIM images of the **(a)** first and **(b)** second centrosome of third instar *cnb* mutant larval neuroblasts, expressing YFP::Cnb^T4A,T9A,S82A^ (white; middle row, green; bottom row), a mutant version of Cnb in which all three consensus phosphorylation sites for Polo were substituted by alanine ^18^. Brains were stained for Asl (top row: white; bottom row: magenta). Orange “M” and yellow “D” stand for mother and daughter centriole respectively. The numbers indicate the Daughter/Mother intensity ration of the representative image. Colored arrowheads and bars underneath the images highlight the degree of Cnb relocalization (see c). **(c)** Graph showing the timeline of Cnb’s relocalization at defined mitotic stages in YFP::Cnb^T4A,T9A,S82A^ expressing *cnb* mutant neuroblasts. The bars show the percentage of neuroblasts containing a single Cnb^+^ centriole (dark blue), a single centriole without Cnb (dark grey), Cnb on both centrioles (transition stage with a Daughter/Mother ratio < 2; light green), predominant Cnb localization on the daughter centriole (strong asymmetry with a Daughter/Mother ratio between 2 and 10; light blue) or in which Cnb is completely shifted to the daughter centriole (complete asymmetry with a Daughter/Mother ratio > 10; light brown). In contrast to wild type Cnb **(d)**, the relocalization of YFP::Cnb^T4A,T9A,S82A^ **(e)** is delayed, which should give rise to two Cnb^+^ interphase centrioles as tested in the following panels. **(f)** Localization of YFP::Cnb^T4A,T9A,S82A^ in *cnb* mutant interphase neuroblasts. **(g)** Quantification of interphase neuroblast phenotype for control (*polo*/+) and Cnb^T4A,T9A,S82A^; *cnb*. These experiments were done twice for YFP::Cnb^T4A,T9A,S82A^; *cnb*. Representative live cell imaging sequence of a **(h)** control neuroblast, expressing wild type YFP::Cnb (green) and **(i)** *cnb* mutant neuroblast expressing YFP::Cnb^T4A,T9A,S82A^ (green). Both samples also co-express the spindle marker UAS-mCherry::Jupiter (white) to visualize microtubules. **(j)** Quantification of centriole splitting phenotype; blue bars represent neuroblasts retaining a single Cnb^+^ centriole on the apical centrosome. Orange bars represent neuroblasts generating two Cnb^+^ centrioles in early interphase. Live cell imaging experiments were repeated 4 times independently. Cell cycle stages are indicated with colored boxes. Time scale is mm:ss. Scale bar is 0.3 μm in a, b, f and 5μm (top row) or 2μm (bottom row) in h and i.

The establishment of molecularly distinct centrioles during mitosis could determine centrosome asymmetry in the following interphase. If so, we would expect that in cases with a strong Cnb relocalization delay, as shown for Cnb phosphomutants, we should find two Cnb^+^ interphase centrosomes (Fig. 3d, e). Indeed, in contrast to wild type, control (*polo*/+) or hypomorphic *polo* mutant neuroblasts, ∼ 75% of *cnb* mutant neuroblasts expressing YFP::Cnb^T4A,T9A,S82A^ contain two Cnb^+^ interphase centrosomes (Fig. 3f, g; Supplementary Fig.2e-g). To more directly visualize the origin of the two Cnb^+^ interphase centrosomes, we imaged *cnb* mutants expressing YFP::Cnb^T4A,T9A,S82A^ live. Unfortunately, we could not obtain reliable live 3D-SIM data and our spinning disc live cell imaging setup cannot resolve individual centrioles during mitosis. However, we reasoned that we could visualize two Cnb^+^ centrioles when the mother and daughter centrioles separate from each other at the end of telophase (Supplementary Fig.1a). Indeed, in contrast to wild type, retaining a single Cnb^+^ centriole on the apical cortex in most neuroblasts (75%; n = 20), we found two separating Cnb^+^ centrioles in most *cnb* mutants expressing YFP::Cnb^T4A,T9A,S82A^ (73%; n = 22; Fig. 3h-j). This data suggests that incomplete relocalization of phosphomutant Cnb from the mother to the daughter centriole during mitosis gives rise to two Cnb^+^ centrioles by the end of telophase. In contrast to wild type, this incomplete phosphomutant Cnb relocalization will result in neuroblasts reentering the next mitosis with Cnb+ on both centrosomes. Taken together, we conclude that Polo dependent phosphorylation of Cnb is necessary for Cnb’s timely relocalization from the mother to the daughter centriole, and that the establishment of molecularly distinct centrioles during mitosis determines subsequent molecular interphase asymmetry.

### Polo becomes enriched on the daughter centriole whereas Plp remains localized on the mother centriole

Having implicated Polo in Cnb’s mother – daughter centriole relocalization we then analyzed the localization of Polo (Polo::GFP) and Plp (Plp::EGFP). The latter has previously been shown to be involved in centrosome asymmetry establishment ^21^. Both Polo and Plp were GFP-tagged at the endogenous locus (^30^ and methods). In early prophase neuroblasts, Polo was localized on the existing centriole on both centrosomes (Fig. 4a, b & ^19^). Subsequently, Polo intensity increased on the forming daughter centriole and its asymmetric localization peaked in metaphase/anaphase. Interestingly, the apical centrosome showed a less pronounced asymmetric distribution in prometaphase compared to the basal centrosome, which could reflect differences in the relocalization mechanism (Fig. 4a-c).

**Fig. 4:**
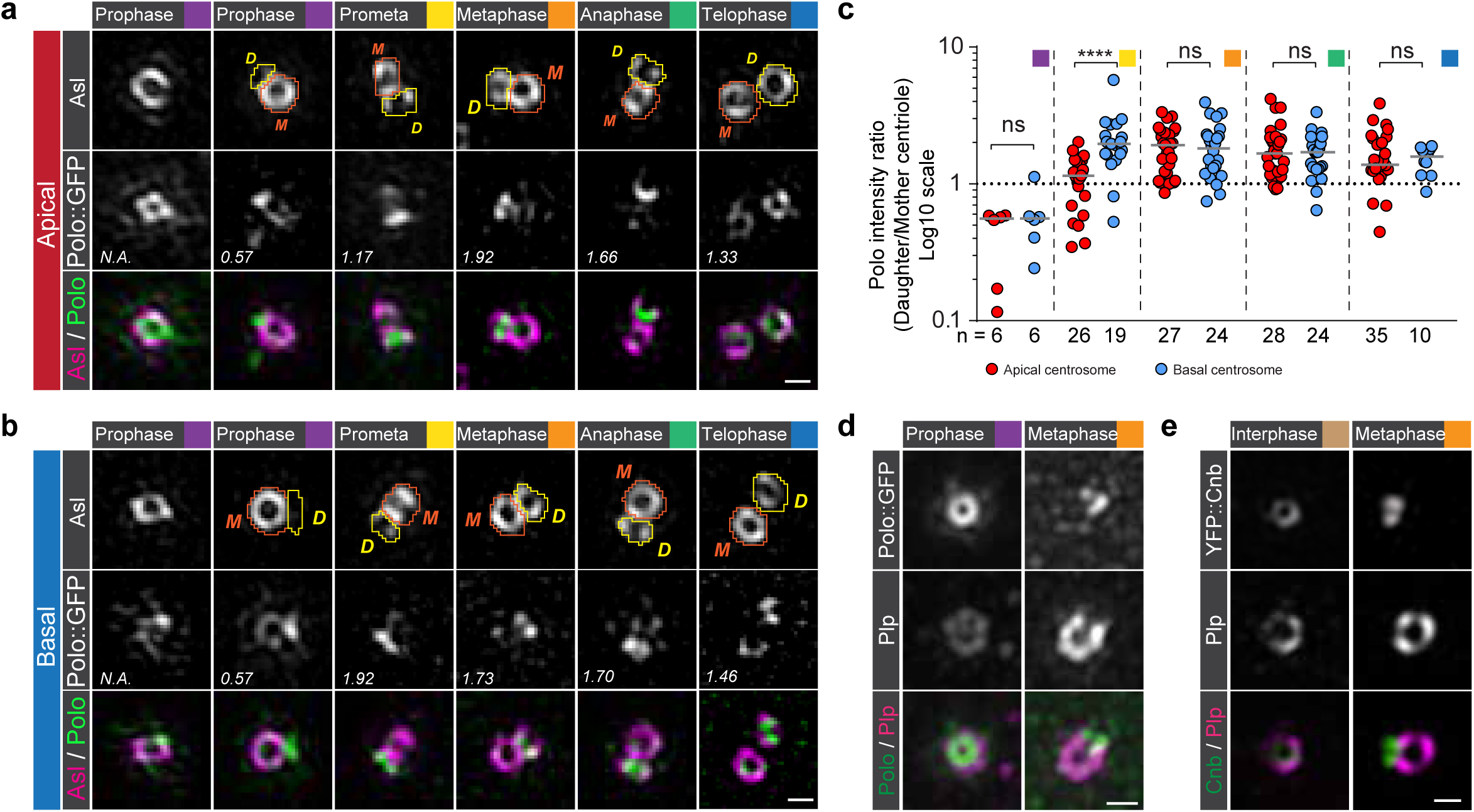
Polo and Cnb separate from Plp in mitosis. Representative 3D-SIM images of **(a)** apical or **(b)** basal third instar larval neuroblast centrioles, expressing Polo::GFP (middle row; green in merge). Centriole contours were drawn based on Asl signal (orange and yellow lines for mother and daughter centriole respectively) and used to measure Polo::GFP (and Asl; not shown) intensities. The numbers represent total Polo intensity ratios (Daughter/Mother centriole) in the shown image. Polo asymmetry ratios for the apical (red dots) and the basal (blue dots) centrosome are plotted in **(c)** from three independent experiments. Medians are shown with a grey horizontal line. Prophase: apical versus basal; p=0.6991. Prometaphase: apical versus basal; p=5.688×10^−6^. Metaphase: apical versus basal; p=0.9329. Anaphase: apical versus basal; p=0.8628. Telophase: apical versus basal: p=0.8614. Representative interpolated images of apical interphase/early prophase and late metaphase/early anaphase centrosomes, expressing **(d)** Polo::GFP (green in merge) or **(e)** YFP::Cnb (green in merge) and stained for Plp (magenta in merge). These experiments were performed three times independently for Polo::GFP and once for YFP::Cnb. Cell cycle stages are indicated with colored boxes. Scale bar is 0.3 μm.

In contrast to Polo and Cnb, Plp predominantly remained localized on the mother centriole on both centrosomes, although it increased also on the daughter centriole in late mitosis (Supplementary Fig.3). Co-imaging Polo together with Plp, and Cnb with Plp showed that Polo and Cnb separated from Plp in metaphase and anaphase (Fig. 4d, e). These data suggest that similar to Cnb on the apical centrosome, Polo is changing its localization from the mother to the daughter centriole during mitosis. However, in contrast to Cnb, Polo’s relocalization dynamics appear similar on both centrosomes. Plp remains enriched on the mother centriole on both the basal and apical centrosome.

### Polo’s relocalization to the daughter centriole depends on Wdr62 and Cnb, with Polo and Cnb co-depending on each other

We next asked how asymmetric Polo localization establishment is regulated. To this end, we analyzed Polo localization in neuroblasts depleted for Cnb (*cnb* RNAi) and Wdr62 (*wdr62* mutants). Wdr62 is implicated in primary microcephaly ^31,32^, and both Cnb and Wdr62 are necessary for MTOC asymmetry by regulating Polo’s and Plp’s centrosomal localization in interphase neuroblasts ^18,19^. Lack of Cnb or Wdr62 did not compromise the gradual loading of Asl onto the newly formed centriole in mitotic neuroblasts and Plp localization was still highly asymmetric in favor of the mother centriole (data not shown). However, the asymmetric centriolar localization of Polo, especially in prometaphase to anaphase neuroblasts, was significantly perturbed in the absence of Cnb and Wdr62 (Fig. 5a-c). Lack of Cnb - but not Wdr62 - also compromised Polo’s asymmetric localization in telophase, suggesting a preferential requirement for Wdr62 in metaphase and anaphase.

**Fig. 5:**
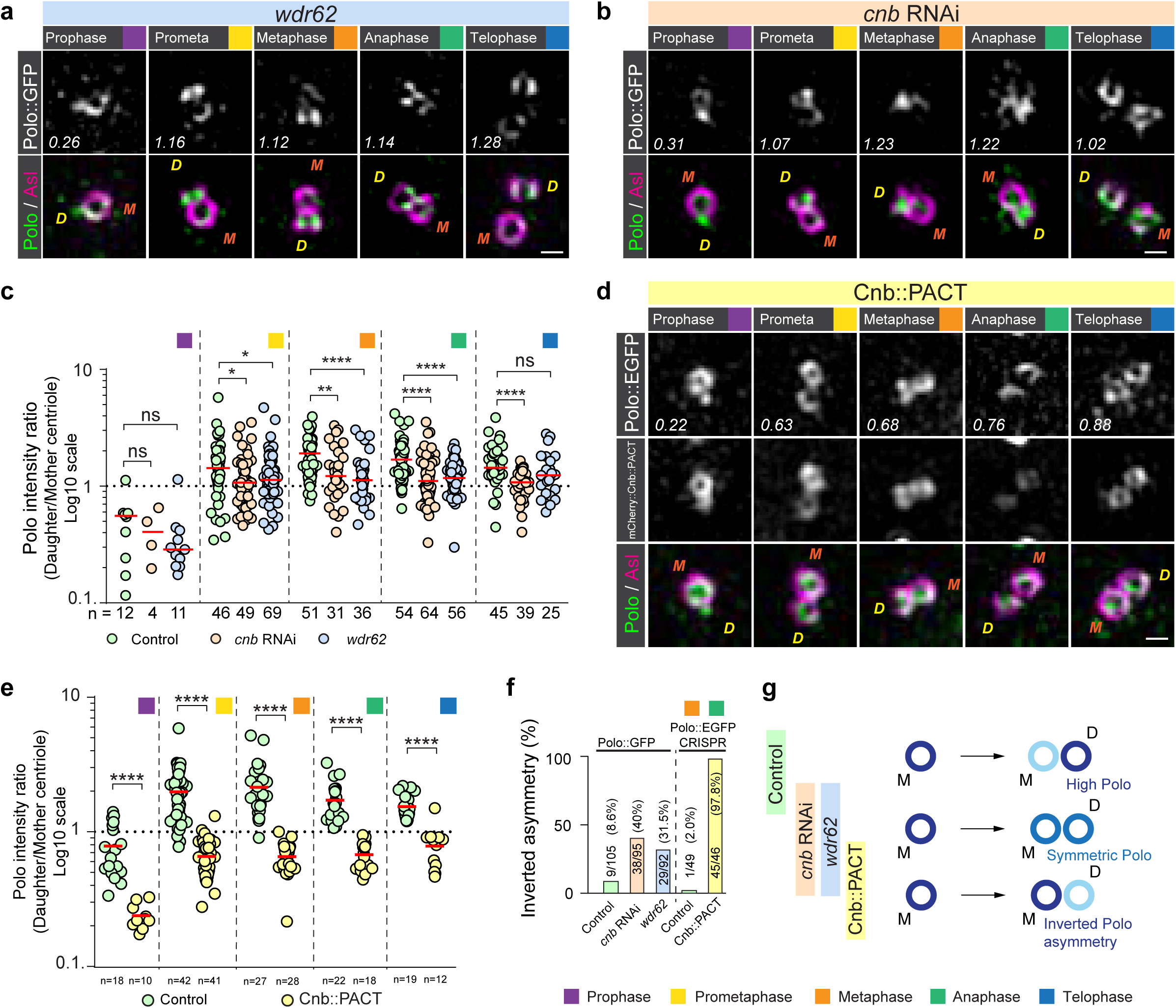
Polo relocalization from the mother to the daughter centriole during mitosis depends on Cnb and Wdr62. Representative 3D-SIM images of third instar larval neuroblasts mutant for **(a)** *wdr62* or **(b)** expressing RNAi against Cnb (*cnb* RNAi). In both conditions, Polo::GFP (green in merge) expressing neuroblasts were stained for Asl (magenta in merge). For all panels, orange “M” and yellow “D” represent mother and daughter centriole respectively. Polo intensity ratios (Daughter/Mother centriole) are shown in the representative images and plotted in **(c)** for control (wild type background; green dots), *cnb* RNAi (beige dots) and *wdr62* mutants (blue dots). Since apical and basal centrosomes could not be distinguished in *cnb* RNAi and *wdr62* mutants, measurements from these conditions were compared to the pooled (apical and basal) control Polo measurements (replotted from Fig. 4c). These experiments were performed three times independently for wild type control and *cnb* RNAi, and six times for *wdr62*. Medians are shown in red. Prophase: wild type control versus *cnb* RNAi; p=0.6835. wild type control versus *wdr62*; p=0.1179. Prometaphase: wild type control versus *cnb* RNAi; p=0.0318. wild type control versus *wdr62*; p=0.0439. Metaphase: wild type control versus *cnb* RNAi; p=0.0040; wild type control versus *wdr62*; p=8.496×10^−5^. Anaphase: wild type control versus *cnb* RNAi; p=4.19×10^−6^. wild type control versus *wdr62*; p=1.79×10^−6^. Telophase: wild type control versus *cnb* RNAi; p=1.17×10^−6^. wild type control versus *wdr62*; p=0.0524. **(d)** Representative 3D-SIM images of third instar larval neuroblast centrosomes, expressing Polo::EGFP generated by CRISPR/Cas9 technology (white, top row; green bottom row) and mCherry::Cnb::PACT (white in middle row), stained for Asl (magenta in merge; bottom row). Polo intensity ratios (Daughter/Mother centriole) are shown in the representative images and plotted in **(e)** for control (wild type background; green dots) and mCherry::Cnb::PACT expressing neuroblasts (yellow dots). Medians are shown in red. This experiment was performed two times independently for wild type and mCherry::Cnb::PACT expressing neuroblast in parallel. Prophase: wild type control versus mCherry::Cnb::PACT; p=1.524 x10^−7^. Prometaphase: wild type control versus mCherry::Cnb::PACT; p<1.0 x10^−15^. Metaphase: wild type control versus mCherry::Cnb::PACT; p=2.0 x10^−15^. Anaphase: wild type control versus mCherry::Cnb::PACT; p=1.764×10^−11^. Telophase: wild type control versus mCherry::Cnb::PACT; p=3.854×10^−6^. The percentage of metaphase and anaphase centrosomes with inverted Polo asymmetry (Daughter/Mother ratio <1) are plotted in **(f)**. **(g)** Summary of phenotypes; efficient relocalization of Polo from the mother (M) to the daughter (D) centriole is prevented in neuroblasts devoid of Wdr62 or Cnb, or with mislocalized Cnb. Cell cycle stages are indicated with colored boxes. Scale bar is 0.3 μm.

Our *polo* mutant, Cnb phosphomutant and Cnb RNAi data are consistent with previous studies, indicating a co-dependency of Polo and Cnb ^18,21^. To test whether Cnb mislocalization is sufficient to prevent Polo relocalization to the daughter centriole, we expressed mCherry::Cnb::PACT (see Methods) together with Polo::EGFP (tagged endogenously, using CRISPR/Cas9 technology; see methods). Since our 3D-SIM data showed Plp to be predominantly associated with the mother centriole, we reasoned that tethering Cnb to the mother centriole with Plp’s PACT domain ^33^ would compromise the establishment of a Cnb^−^ mother and Cnb^+^ daughter centriole. We speculated that Cnb’s localization would remain enriched on the mother centriole or at least become near symmetrically localized. Indeed, our 3D-SIM experiments revealed that mCherry::Cnb::PACT or YFP::Cnb::PACT ^18^ failed to properly relocalize from the mother to the daughter centriole and remained associated with the mother centriole (Fig. 5d & Supplementary Fig.4a, b). Tethering the PACT domain to Cnb prevented the establishment of a high daughter/mother centriole Polo asymmetry. Polo::EGFP was either localized symmetrically (with equal amounts on both the mother or daughter centriole) or, as observed in most cases, inverted asymmetrically (with higher Polo::EGFP amounts on the mother centriole) (Fig. 5d-e). Taken together, loss or mislocalization of Cnb and depletion of *wdr62* significantly increased the number of centrosomes with inverted Polo asymmetry ratios (wild type control: 8.6% and 2%, respectively; *cnb* RNAi: 40%; *wdr62*: 31.5%; Cnb::PACT: 97.8%; Fig. 5f, g). We conclude that Wdr62 and Cnb are necessary to establish ‘low-Polo’ mother and ‘high-Polo’ daughter centriole containing centrosomes during mitosis. Furthermore, Polo and Cnb both co-depend on each other to correctly establish this centriolar asymmetry.

### Disrupting centriolar asymmetry impacts biased MTOC activity in interphase and spindle orientation in metaphase

Next, we set out to investigate the significance of centriole asymmetry establishment by preventing the relocalization of Cnb and Polo from the mother to the daughter centriole using the PACT domain (see above). It was previously shown that expression of YFP::Cnb::PACT in neuroblasts converted the inactive mother interphase centrosome into an active MTOC, resulting in the presence of two active interphase MTOCs ^18^ (Supplementary Fig.4c, Movie 1&2). However, the underlying mechanisms have not been further investigated. We hypothesized that fusing Cnb with the PACT domain affects the correct establishment of molecular centrosome asymmetry during mitosis, manifested in symmetric MTOC activity in the subsequent interphase. To test this hypothesis, we developed a nanobody trapping experiment, using the anti-GFP single domain antibody fragment (vhhGFP4) ^34,35^ fused to the PACT domain of Plp ^33^ to predominantly trap GFP- or YFP-tagged proteins on the mother centriole (Supplementary Fig.5a-c). Expressing PACT::vhhGFP4 in neuroblasts together with YFP::Cnb mimics the YFP::Cnb::PACT phenotype; almost 93% (n = 69) of neuroblasts expressing PACT::vhhGFP4 together with YFP::Cnb showed two active interphase MTOCs (YFP::Cnb expression only: no MTOC gain of function observed; n = 16; Supplementary Fig.5d, E; Movie 3). Conversely, trapping Asl::GFP with PACT::vhhGFP4 on the mother centriole did not cause a MTOC phenotype in 83% of neuroblasts (n = 104; Supplementary Fig.5f, g; Movie 4).

Having validated the nanobody tool, we next co-expressed a GFP-tagged version of Polo – either a published GFP::Polo transgene ^36^ or our endogenously tagged CRISPR Polo::EGFP line – with PACT::vhhGFP4. 3D-SIM data revealed that under these experimental conditions, Polo::EGFP was strongly localized to the mother centriole in prophase. Subsequently, Polo::EGFP was symmetrically localized between mother and daughter centriole from prometaphase onwards (Supplementary Fig.6a, b). Nanobody-mediated trapping of Polo on the mother centriole also induced the formation of two active interphase MTOCs (GFP::Polo transgene: 84%; n = 31. Polo::EGFP CRISPR line: 72%; n = 82) (Supplementary Fig.5h, i, Supplementary Fig.6c-e & Movie 5-7). Although cell cycle progression was not affected in these neuroblasts, we measured a significant misorientation of the mitotic spindle in early metaphase (Supplementary Fig.6f-g, i). However, similar to *bld10* mutant neuroblasts, displaying two active interphase MTOCs also ^20^, mitotic spindles realigned along the apical-basal polarity axis, ensuring normal asymmetric cell divisions along a conserved axis between successive mitoses (Supplementary Fig.6h, j). 3D-SIM imaging also revealed that in Polo::EGFP & vhhGFP4::PACT expressing neuroblast, both interphase centrosomes (now containing one centriole each) contain high levels of centriolar and diffuse PCM Polo, consistent with our recent observation for the apical interphase wild type centrosome ^19^ (Supplementary Fig.6k, l). Based on these experiments we conclude that preventing the normal establishment of Cnb and Polo asymmetry using the PACT domain perturbs biased MTOC activity in interphase.

### Optogenetically induced Polo and Cnb trapping during mitosis affects MTOC activity in the subsequent interphase

Based on these nanobody results, we reasoned that trapping Polo and Cnb on the mother centriole at defined cell cycle stages should allow us to test more specifically whether the establishment of Polo and Cnb asymmetry during mitosis has an impact on MTOC activity in the subsequent interphase. To test this hypothesis, we implemented the optogenetic system iLID ^37^ by generating transgenic flies containing the iLID cassette (containing the *Avena Sativa’s* LOV domain) fused with the PACT domain (*UAS-iLID::PACT::HA; UAS-iLID::PACT::GFP*; see methods). iLID (or SsrA) binds to the small SspB domain under blue light exposure ^37^. To test this system in fly neuroblasts, we expressed cytoplasmic SspB::mCherry together with iLID::PACT::GFP and exposed entire larval brains first to yellow (561nm) light, followed by simultaneous blue and yellow light (488 and 561nm) exposure, before switching back to only 561nm; each exposure period lasted 5 minutes. Blue light exposure was sufficient to induce the recruitment of cytoplasmic SspB::mCherry to neuroblast centrioles containing iLID::PACT::GFP within 15 seconds. This behavior is strictly blue-light dependent as imaging with 561nm alone is not sufficient to recruit SspB::mCherry to centrioles and SspB::mCherry relocalized to the cytoplasm within 100s after blue light exposure was shut off (Supplementary Fig.7a).

Next, we generated *SspB::EGFP::Polo* and *SspB::mDendra2::Cnb* flies using CRISPR/Cas9. We reared embryos, expressing iLID::PACT::HA under the control of the neuroblast specific *worGal4* driver together with SspB::EGFP::Polo or SspB::mDendra::Cnb in the dark for 4 days before exposing third instar larval neuroblasts in intact brains to blue light at different cell cycle stages for 10-20 minutes. Subsequently, we monitored MT dynamics using mCherry::Jupiter for ∼ 90 minutes without blue light exposure. If the dynamic relocalization of Polo and Cnb during mitosis is important for the correct MTOC establishment in the subsequent interphase (interphase 2), we would expect that light-dependent manipulation of Cnb and Polo localization would mimic the nanobody phenotype, resulting in two active MTOCs in interphase 2. Indeed, many neuroblasts, exposed to blue light from late interphase 1 or prophase 1 onward, showed two active MTOCs in the following interphase 2. However, continued blue light exposure during interphase – early interphase in particular - also disrupted MTOC asymmetry in late interphase just prior to mitotic entry (Fig. 6a-c). Overall, ∼ 55 % of SspB::EGFP::Polo & iLID::PACT::HA and ∼ 47 % of SspB::mDendra2::Cnb & iLID::PACT::GFP expressing neuroblasts, exposed to blue light showed an MTOC phenotype (n = 67 and n = 40, respectively; Fig. 6d, e). We restricted the analysis to neuroblasts showing an MTOC phenotype since the efficiency of optogenetic recruitment is variable making a negative result difficult to interpret. SspB::EGFP::Polo also displayed a more focused and intense localization when co-expressed with iLID::PACT::HA and exposed to blue light, compared to normal SspB::EGFP::Polo localization (Supplementary Fig.7b). These observed phenotypes are strictly blue light dependent as SspB::EGFP::Polo or SspB::mDendra2::Cnb expressed in conjunction with iLID::PACT and imaged with 561nm only, showed predominantly normal MTOC activity (SspB::Polo: 87.8% normal divisions; n = 29; SspB::Cnb: 82.3% normal divisions; n = 36; Fig. 6d, e). Taken together, these experiments suggest that perturbing normal Cnb and Polo relocalization during mitosis disrupts asymmetric MTOC behavior in the following interphase. The data further indicates that neuroblasts are also sensitive to optogenetic manipulation of Cnb and Polo localization during interphase.

**Fig. 6:**
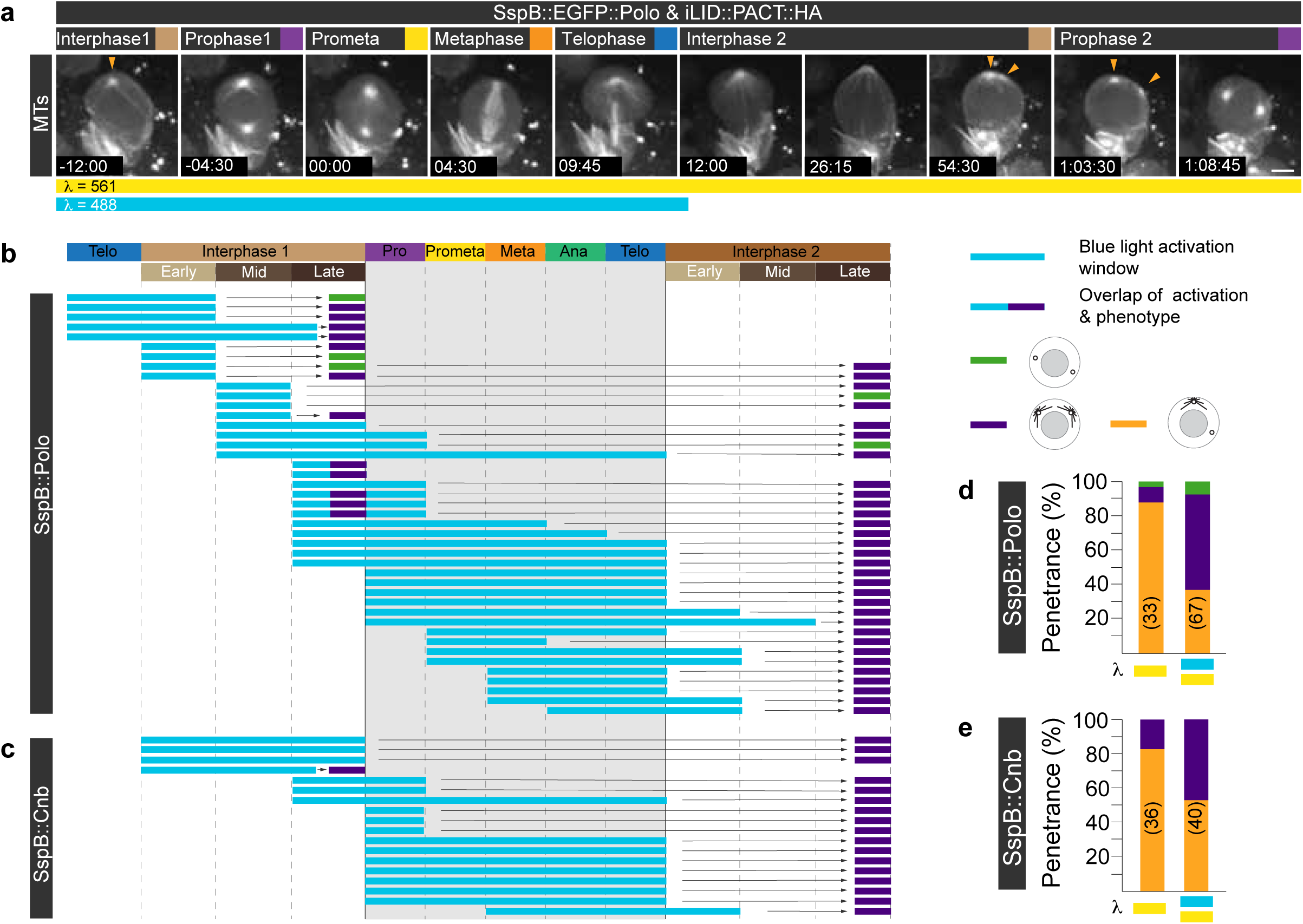
Establishment of centriolar asymmetry during mitosis is required for biased interphase MTOC activity. **(a)** Representative wild type neuroblast expressing SspB::EGFP::Polo (not shown) together with the microtubule marker mCherry::Jupiter (white) and iLID::PACT::HA (not shown). As indicated with the cyan and yellow color bars underneath the image sequence, this neuroblast was exposed to both 488nm and 561nm during the first division but only to 561nm in the second division. Yellow arrowheads indicate two active MTOCs in the interphase 2 and prophase 2. Summary of all optogenetic experiments for **(b)** SspB::EGFP::Polo and **(c)** SspB::mDendra2::Cnb and iLID::PACT::HA expressing neuroblasts. Blue light exposure and resulting phenotype are indicated with the colored bars (see legend on the right). This experiment was repeated more than 3 times independently. Bar graphs representing the phenotypic penetrance (in %) of larvae expressing **(d)** SspB::EGFP::Polo & iLID::PACT::HA or **(e)** SspB::mDendra2::Cnb & iLID::PACT::GFP with (cyan and yellow bar) or without (yellow bar only) blue light exposure. The number of scored divisions are indicated on the bars.

## Discussion

Centrosome asymmetry has previously been described to occur in asymmetrically dividing *Drosophila* neural stem cells (neuroblasts), manifested in biased interphase MTOC activity and asymmetric localization of the centrosomal proteins Cnb, Plp and Polo ^11,18-21^. Here, we have shown that neuroblast centrosomes become intrinsically asymmetric by relocalizing centriolar proteins such as Cnb and Polo from the old mother to the young daughter centriole during mitosis. This establishment of centriolar asymmetry is tightly linked to centriole-to-centrosome/mitotic centriole conversion ^25,27^. In early prophase, Cnb and Polo colocalize on the existing mother centriole of the apical centrosome but from late prometaphase onward, Cnb and Polo are exclusively (in the case of Cnb) or predominantly (in the case of Polo) localized on the daughter centriole. Interestingly, Cnb behaves differently on the basal centrosome: the existing mother centriole does not contain any Cnb, appearing only on the forming daughter centriole in late prophase. On the apical centrosome however, Cnb is often present on the mother and daughter centriole between late prophase and early prometaphase. Mechanistically, the relocalization could entail a direct translocation of Cnb and Polo from the mother to the daughter centriole. This model is partially supported by our FRAP data. However, on the basal centrosome, Cnb is completely absent from the existing mother, and appears only in late prophase on the forming daughter centriole. This suggests a direct recruitment mechanism, which could also apply to the apical centrosome from anaphase onward. We propose a dual mechanism whereby on the apical centrosome Cnb initially directly transfers from the mother to the daughter centriole. From anaphase onward – and from late prophase onward on the basal daughter centriole – Cnb levels increase through direct recruitment (Fig. 6f, g). Cnb is phosphorylated by the mitotic kinase Polo ^18^ and Polo-dependent phosphorylation of Cnb is necessary for its timely relocalization during mitosis, suggesting that Polo regulates the dynamic relocalization of Cnb from the mother to the daughter centriole. Interestingly, our data further suggest that Polo, which also becomes enriched on the daughter centriole during mitosis is co-dependent with Cnb, while also requiring Wdr62. Polo’s involvement in mitotic centriole conversion ^27^ further suggests that the same molecular machinery cooperatively converts a maturing centriole into a centrosome for the next cell cycle while simultaneously providing it with its unique molecular identity (Fig. 7a - c).

**Fig. 7:**
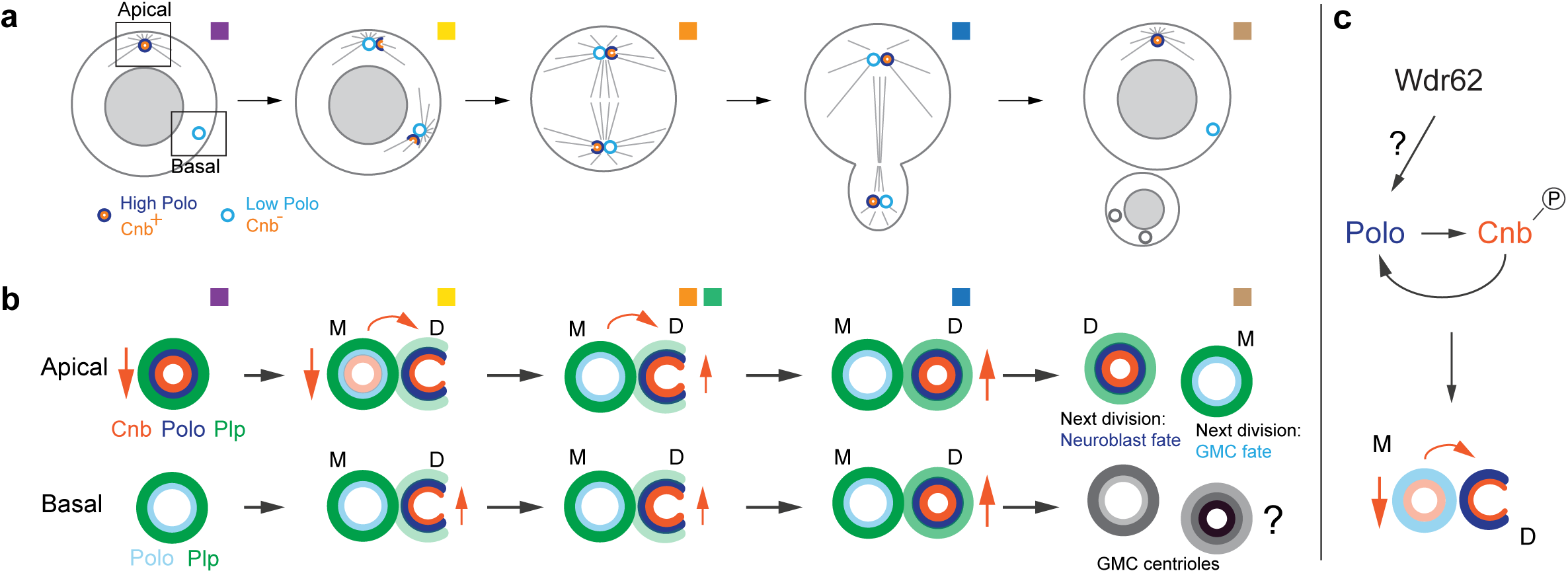
Centrosome asymmetry is primed in mitosis through dynamic Cnb and Polo relocalization. **(a)** Model: Centriolar asymmetry – here shown for Polo (dark and light blue) and Cnb (orange) – occurs during mitosis, coupled to centriole-to-centrosome/mitotic centriole conversion. Polo and Cnb are relocalizing from the existing mother to the newly formed daughter centriole on both the apical and basal centrosome. The ensuing Polo-rich centriole maintains MTOC activity, retaining it in the self-renewed neuroblast. Details for the apical and basal centrosome are shown in **(b)**. Cnb (orange) and Polo (blue) relocalize from the mother to the forming daughter centriole from prophase onwards. The basal centrosome only relocalizes Polo but directly upregulates Cnb on the daughter centriole. Cnb’s relocalization most likely entails both down and upregulation in prophase/prometaphase and upregulation in anaphase/telophase, respectively (vertical orange arrows), as well as direct protein transfer (curved arrows). Plp (green) remains on the mother, potentially increasing in intensity and appearing on the daughter centriole in prometaphase. Centriolar protein relocalization is mostly completed by anaphase. The centriole containing less Plp, gained Cnb and Polo and is destined to be inherited by the self-renewed neuroblast (indicated with ‘neuroblast fate’) in the next division, whereas the centriole containing higher Plp and lower Polo levels is destined to be inherited by the GMC (indicated with ‘GMC fate’). The fate of the basal centrioles and subsequent marker distribution is unknown (represented by grey circles). **(c)** Cnb and Polo co-depend on each other for their relocalization from the mother to the daughter centriole. Wdr62 is necessary for Polo relocalization albeit the molecular mechanism is unclear. Time scale is mm:ss or hh:mm:ss. Scale bar is 5μm.

The mechanisms generating two molecularly distinct centrioles during mitosis seem to directly influence the centrosome’s MTOC activity in interphase; the ‘Cnb^+^, high Polo’ daughter centriole will retain MTOC activity during interphase whereas the ‘Cnb^−^, low Polo’ mother centriole, separates from its daughter in early interphase and becomes inactive ^18-21,38^. This model is in agreement with *bld10* or *plp* mutants, which fail to downregulate Polo from the mother centriole, resulting in the formation of two active interphase MTOCs ^20,21^. It is further supported by our mislocalization data. For example, optogenetic manipulation of Polo and Cnb asymmetry specifically during mitosis impacts MTOC activity in the subsequent interphase. However, we cannot exclude the possibility that MTOC asymmetry is also controlled independently of mitotic centrosome asymmetry establishment since optogenetic interphase manipulations of Polo and Cnb alone can also perturb biased MTOC activity.

Loss of Wdr62 or Cnb also affects asymmetric centriolar Polo localization. Yet, interphase centrosomes lose their activity in these mutants. *wdr62* mutants and *cnb* RNAi neuroblasts both show low Polo levels in interphase ^19^. We thus hypothesize that in addition to an asymmetric distribution, Polo levels must remain at a certain level to maintain interphase MTOC activity; high symmetric Polo results in two active interphase MTOCs whereas low symmetric Polo results in the formation of two inactive centrosomes. Indeed, our optogenetic experiment revealed increased centriolar Polo levels upon blue light induction, suggesting that both Polo levels and distribution influence MTOC activity.

Taken together, the results reported here are consistent with a model, proposing that the establishment of two molecularly distinct centrioles is primed during mitosis, and contributes to biased MTOC activity in the subsequent interphase. Wild type neuroblasts unequally distribute a given pool of Cnb and Polo protein between the two centrioles so that the centriole inheriting high amounts of Cnb and Polo will retain MTOC activity. Furthermore, the dynamic relocalization of Polo and Cnb provides a molecular explanation for why the daughter centriole-containing centrosome remains tethered to the apical neuroblast cortex and is being inherited by the self-renewed neuroblast ^19-21^ (Fig. 7a). It remains to be tested why neuroblasts implemented such a robust machinery to asymmetrically segregate the daughter-containing centriole to the self-renewed neuroblast; more refined molecular and behavioral assays will be necessary to elucidate the developmental and post-developmental consequences of biased centrosome segregation. The tools and findings reported here will be instrumental in targeted perturbations of intrinsic centrosome asymmetry with spatiotemporal precision in defined neuroblast lineages.

Finally, our observations reported here further raise the tantalizing possibility that centriolar proteins also dynamically relocalize in other stem cells, potentially providing a mechanistic explanation for the differences in centriole inheritance across different stem cell systems.

## Acknowledgements

We thank members of the Cabernard lab for helpful discussions. We are grateful to Jordan Raff, Nasser Rusan, Tomer Avidor-Reiss, Cayetano Gonzalez and Chris Doe for flies and antibodies. We would also like to thank the Imaging Core Facility (IMCF) at the Biozentrum for technical support and the Nigg and Affolter labs for providing temporary lab space to E. G. This work was supported by the Swiss National Science Foundation (SNSF; PP00P3_159318), the National Institutes of Health (NIH; 1R01GM126029-01) and start-up funds from the University of Washington. E.G was supported with an EMBO long-term postdoctoral fellowship (ALTF 378-2015). Stocks obtained from the Bloomington Drosophila Stock Center (NIH P40OD018537) and the Vienna Drosophila Resource Center (VDRC) were used in this study.

## Author contributions

This study was conceived by A.R.N, P.S, and C.C.

E.G and A.R.N performed most of the 3D-SIM experiments with help from P.S. E.G performed all the nanobody experiments. A. M generated the PACT::vhhGFP4 and iLID::PACT constructs. N.H generated the SspB::mDendra::Cnb and SspB::EGFP::Polo constructs and performed the majority of the optogenetic experiments. T.P wrote custom-made Matlab codes and helped with data analysis. D.S.G generated the Plp CRISPR line and A.F helped with 3D-SIM imaging. C.C generated and analyzed Cnb phosphomutant live cell imaging data, FRAP and optogenetic data. E.G, A.R.N and C.C wrote the paper.

## Materials & Correspondence

Material requests and other inquiries should be directed to.

## Supplementary Figure legends

**Supplementary Fig.1.: Centriole duplication completes during mitosis in larval neuroblasts**

**(a)** Current model of centrosome asymmetry in neuroblast. The Cnb^+^ apical daughter centrosome is active throughout interphase and constantly nucleates a robust microtubule array, maintaining its position at the apical neuroblast cortex (blue crescent). The Cnb^−^ basal mother centrosome is inactive during interphase, diffusing through the cytoplasm until it regains MTOC activity in prophase. At this point, the Cnb^−^ centrosome reached the basal side of the neuroblast and starts to reaccumulate Cnb during mitosis. The daughter centrosome is retained by the neuroblast and the mother centrosome is inherited by the differentiating GMC. Asymmetric centrosomes split in early interphase. **(b)** Representative 3D-SIM images of neuroblasts expressing the pericentriolar marker Cnn::GFP stained for α-Tubulin, labelling microtubules (MTs; green). The morphology of the microtubule array and cell shape were used to define neuroblast cell cycle stages. **(c)** Neuroblast centrosomes are inherently asymmetric in interphase but when neuroblast centrioles duplicate and acquire a unique molecular identity (indicated by arrow and color switch) is unknown. **(d)** Representative interpolated 3D-SIM images of third instar larval neuroblast centrosomes, expressing Sas-6::GFP (top row; white. Green in merge) and stained for Asl (middle row; white. Merged channels; red). The yellow arrowhead highlights the cartwheel of the forming centriole. Cartwheel duplication can be observed at the telophase/interphase transition, concomitantly with centrosome separation (blue arrowhead). Cell cycle stages are indicated with colored boxes. Scale bar is 3 μm in (b) and 0.3 μm in (d).

**Supplementary Fig.2.: Cnb’s relocalization is controlled by Polo-dependent phosphorylation**

Representative 3D-SIM images of the **(a)** first and **(b)** second centrosome of a *polo* mutant (*polo^1^/polo^16-1^*) third instar larval neuroblast expressing YFP::Cnb (white; middle row, green; bottom row) and stained for Asl (white; top row, magenta; bottom row). *polo* mutant neuroblasts show a loss of MTOC activity during interphase, which randomizes centrosome positioning and distribution. Since we cannot distinguish between the ‘apical’ and ‘basal’ centrosomes anymore we refer to centrosome 1 and 2 instead. Colored arrowheads and bars underneath the images highlight the degree of Cnb relocalization. Graphs showing the timeline of Cnb’s relocalization at defined mitotic stages in **(c)** control (*polo/+*) and **(d)** *polo* mutant (*polo^1^/polo^16-1^*) neuroblasts. The bars show the percentage of neuroblasts containing a single Cnb^+^ centriole (dark blue), a single centriole without Cnb (dark grey), Cnb on both centrioles (transition stage with a Daughter/Mother ratio < 2; light green), predominant Cnb localization on the daughter centriole (strong asymmetry with a Daughter/Mother ratio between 2 and 10; light blue) or in which Cnb is completely shifted to the daughter centriole (complete asymmetry with a Daughter/Mother ratio > 10; light brown). The localization of YFP::Cnb is shown in **(e)** control and **(f)** *polo* mutant (*polo^1^/polo^16-1^*) interphase neuroblasts. The quantification is displayed in **(g).** This experiment was done three times. Scale bar is 0.3 μm

**Supplementary Fig.3.: Plp remains enriched on the mother centriole during mitosis**

Representative 3D-SIM images of **(a)** apical and **(b)** basal third instar larval neuroblast centrosomes, expressing Plp::EGFP (white in middle row, green in merge), co-stained with Asl (white on top, magenta in merge). Orange and yellow shapes represent mother (M) and daughter (D) centrioles respectively, based on Asl intensity. The number represents total Plp intensity ratios (Daughter/Mother centriole) in the shown image. Plp asymmetry ratios for the apical (red dots) and the basal (blue dots) centrosome are plotted in **(c)** from three independent experiments. Medians are shown in dark grey. Prometaphase: apical versus basal; p=0.3856. Metaphase: apical versus basal; p=0.2234. Anaphase: apical versus basal; p=0.3583. Telophase: apical versus basal; p=0.1844. Plp remains localized on the mother centriole on both centrosomes and enriches on the daughter centriole over time. Scale bar is 0.3 μm. Colored boxes indicate cell cycle stages.

**Supplementary Fig.4.: YFP::Cnb::PACT expression impairs complete Cnb relocalization from the mother to the daughter centriole, affecting interphase MTOC activity**

**(a)** Representative 3D-SIM images of third instar larval neuroblast centrosomes, expressing YFP::Cnb::PACT (white in the second row, green in the merge) and stained for Asl (white in the first row, magenta in the merge). The number represents total YFP::Cnb::PACT intensity ratios (Daughter/Mother centriole) in the shown image. YFP::Cnb::PACT intensity ratios (Daughter/Mother centriole) are plotted in **(b)**. **(c)** Representative live cell imaging series from a neuroblast, recorded in the intact brain, expressing the microtubule marker mCherry::Jupiter (MTs, first row) and YFP::Cnb::PACT (second row). Red and blue squares represent apical and basal centrosome respectively. 3D-SIM and live imaging experiments were performed two times each. “00:00” corresponds to the telophase of the first division. Cell cycle stages are indicated with colored boxes. Yellow “D” and orange “M” refer to Daughter and Mother centrioles based on Asl intensity. Timestamps are shown in hh:mm and scale bar is 0.3μm (a) and 3 μm (c).

**Supplementary Fig.5.: Perturbing centriolar asymmetry by tethering the GFP-trapping nanobody to the mother centriole**

**(a)** To test the function of centriolar asymmetry establishment, the relocalization of Polo and Cnb needs to be perturbed. **(b)** Nanobody technology was used to prevent the centrosome asymmetry switch for selected proteins of interest. The vhhGFP4 nanobody specifically traps GFP or YFP tagged proteins. By tethering the nanobody preferentially to the mother centriole - using Plp’s PACT domain **(c)**, we can perturb the relocalization of GFP or YFP tagged centrosomal proteins. Crossed-out arrows illustrate a lack of centriolar protein relocalization (shown for Polo; blue). Representative live cell image series from intact brains for neuroblasts expressing the microtubule marker mCherry::Jupiter (first row) and PACT::VhhGFP4 together with **(d)** YFP::Cnb, **(f)** Asl::GFP and **(h)** GFP::Polo transgene (genomic rescue construct; see methods). MTOC phenotype quantifications are shown for **(e)** YFP::Cnb, **(g)** Asl::GFP and **(i)** GFP::Polo (blue; wild type-like asymmetry, dark brown; loss of MTOC activity, light brown; gain of MTOC activity). “00:00” corresponds to telophase of the first division. Cell cycle stages are indicated with colored boxes. The data presented here were obtained from two, four and three independent experiments for YFP::Cnb, Asl::GFP and GFP::Polo respectively. Timestamps are hh:mm and scale bar is 3μm.

**Supplementary Fig.6.: Establishment of centriolar asymmetry is required for biased interphase MTOC activity and centrosome positioning.**

**(a)** Representative 3D-SIM images of third instar larval neuroblast centrosomes, expressing Polo::EGFP (generated by CRISPR/Cas9) and the nanobody construct PACT::vhhGFP4. Polo::EGFP (middle: white; merge: green) expressing neuroblasts were stained for Asl (white; top row, magenta in the merge). Polo intensity ratios (Daughter/Mother centriole) are plotted in **(b)** for control (green dots) and PACT::vhhGFP4 (purple dots). These experiments were performed two times independently in parallel for both genotypes. Medians are shown in red. Prophase: Control versus PACT::vhhGFP4; p=3.11×10^−4^. Prometaphase: Control versus PACT::vhhGFP4; p=3.49×10^−6^. Metaphase: Control versus PACT::vhhGFP4; p=0.0222. Anaphase: Control versus PACT::vhhGFP4; p=6.28×10^−5^. Telophase: Control versus PACT::vhhGFP4; p=0.0077. **(c)** Representative live cell imaging time series of a dividing control (Polo::EGFP, worGal4, UAS-mCherry::Jupiter) and **(d)** PACT::vhhGFP4 expressing (Polo::EGFP, worGal4, UAS-mCherry::Jupiter & PACT::vhhGFP4) neuroblast. The microtubule marker (MTs, first row) and Polo::EGFP (second row) are shown for two consecutive mitoses. Microtubule intensity of the apical (red line and square) and basal (blue line and square) MTOC are plotted below. “00:00” corresponds to the telophase of the first division. **(e)**. Bar graph showing the quantification of the MTOC phenotype in interphase (blue; wild type-like asymmetry, dark brown; loss of MTOC activity, light brown; gain of MTOC activity). Cell cycle length is shown in **(f)**. The cell cycle length in PACT::vhhGFP4 (purple dots) is not significantly different from the control (green dots); p=9727. Medians are shown in red. **(g)** and **(i)** represent the spindle rotation between NEBD and anaphase. Medians are displayed in dark colors (green; control. Purple; vhhGFP4 expressing neuroblasts) and the maximum rotation in light colors. Division orientation between consecutive mitoses shown for control **(h)** and PACT::vhhGFP4 **(j)**. **(k)** and **(l)** are representative 3D-SIM images of interphase centrosomes for control and PACT::vhhGFP4 expressing neuroblasts, respectively. The trapping of Polo::EGFP with PACT::vhhGFP4 induces two identical apical-like (in respect to MTOC activity and Polo localization) centrosomes with a strong centriolar and PCM signal. The data presented for the live imaging here were obtained from five independent experiments. Cell cycle stages are indicated with colored boxes. Yellow “D” and orange “M” refer to Daughter and Mother centrioles based on Asl intensity. Timestamps are shown in hh:mm and scale bar is 0.3μm (a, k, l) and 3 μm (c, d), respectively.

**Supplementary Fig.7.: optogenetic protein relocalization is efficient on third instar larval neuroblast centrosomes**

**(a)** Representative time-lapse frames of a third instar neuroblast – imaged in an intact brain – expressing SspB::mCherry (second and third row; grey) together with iLID::PACT::GFP (cyan; top row). Light exposure regime is indicated on the top. Orange brackets and red arrowheads highlight the apical neuroblast centrosome. An unrelated mCherry particle is highlighted with the green arrowhead. Intensity ratios, displaying the ratio of centrosomal/cytoplasmic SspB::mCherry are shown below; SspB::mCherry intensity was measured along a 12-pixel wide line covering the centriole and normalized against cytoplasmic mCherry levels. Note that SspB::mCherry relocalizes from the cytoplasm to the apical centrosome within 5 seconds and relocalized back into the cytoplasm within ∼ 2 minutes. **(b)** Representative Prophase time-lapse frames of third instar larval neuroblasts expressing SspB::EGFP::Polo alone (control; left) or in conjunction with iLID::PACT::HA (right). SspB::EGFP::Polo (middle row; white) appears enriched and more focused in the presence of iLID::PACT::HA and after blue light exposure. Intensity ratios, displaying the ratio of centrosomal/cytoplasmic SspB::EGFP::Polo are shown below; SspB::EGFP::Polo intensity was measured along a 12-pixel wide line covering the centriole and normalized against cytoplasmic EGFP levels. Time scale is mm:ss. Scale bar is 5μm.

## Description of Additional Supplementary Files

**File Name: Supplementary Movie 1**

**Description:** Wild type neuroblasts expressing YFP::Cnb.

Wild type control larval neuroblast expressing the centriolar marker YFP::Cnb (green) and the microtubule marker UAS-mCherry::Jupiter (white), driven by the neuroblast-specific worGal4 transgene. Note that the daughter centriole (Cnb^+^) remains active and anchored to the apical cortex throughout interphase. The second centrosome matures in prophase (00:39) after it reached the basal side of the cell. “00:00” corresponds to telophase. Time scale is hh:mm and the scale bar is 3μm.

**File Name: Supplementary Movie 2**

**Description:** Wild type neuroblast expressing YFP::Cnb::PACT.

Larval neuroblast expressing YFP::Cnb::PACT (green) and the microtubule marker UAS-mCherry::Jupiter (white), driven by the neuroblast-specific worGal4 transgene. Note that YFP::Cnb::PACT is present on both centrioles. Both centrosomes remain active and anchored to the apical cortex throughout interphase. Centrioles split in prophase (00:39) accompanied by a large spindle rotation (00:42 - 00:45), resulting in normal asymmetric cell division. “00:00” corresponds to telophase. Time scale is hh:mm and the scale bar is 3μm.

**File Name: Supplementary Movie 3**

**Description:** Neuroblast expressing YFP::Cnb together with centriole tethered PACT::vhhGFP4.

Larval neuroblast expressing the centriolar marker YFP::Cnb (green), the microtubule marker UAS-mCherry::Jupiter (white) and the PACT::vhhGFP4 nanobody; both UAS lines are driven by the neuroblast-specific worGal4 transgene. The PACT domain confines the nanobody predominantly to the mother centriole. Both centrosomes remain active and anchored to the apical cortex throughout interphase. Centrosome splitting occurs a few minutes before mitosis (00:36). “00:00” corresponds to telophase. Time scale is hh:mm and the scale bar is 3μm.

**File Name: Supplementary Movie 4**

**Description:** Neuroblast expressing Asl::GFP together with centriole tethered PACT::vhhGFP4.

Larval neuroblast expressing the centriolar marker Asl::GFP (green), the microtubule marker UAS-mCherry::Jupiter (white) and the PACT::vhhGFP4 nanobody; both UAS lines are driven by the neuroblast-specific worGal4 transgene. Similar to the wild type control, the daughter centriole remains active and anchored to the apical cortex throughout interphase. The mother centriole sheds its MTOC activity and moves away in early interphase (00:15). At mitotic entry (00:45), the mother centriole matures after it reached the basal side of the cell. “00:00” corresponds to telophase. Time scale is hh:mm and the scale bar is 3μm.

**File Name: Supplementary Movie 5**

**Description:** Wild type control neuroblast expressing Polo::EGFP.

Wild type control larval neuroblast expressing Polo::EGFP (green) engineered by CRISPR/Cas9 technology and the microtubule marker mCherry::Jupiter (white). Note that the daughter centriole remains active and anchored to the apical cortex throughout interphase. The mother centriole matures at 00:42 after it reached the basal cell cortex. “00:00” corresponds to telophase. Time scale is hh:mm and the scale bar is 3μm.

**File Name: Supplementary Movie 6**

**Description:** Neuroblast expressing Polo::EGFP together with centriole tethered PACT::vhhGFP4. Larval neuroblast expressing Polo::EGFP (green) engineered by CRISPR/Cas9 technology, the microtubule marker mCherry::Jupiter (white) and the PACT::vhhGFP4 nanobody; both UAS lines are driven by the neuroblast-specific worGal4 transgene. Both MTOCs remain active and anchored to the apical cortex throughout interphase. Centrioles split only 6 minutes before mitosis starts (00:36). The mitotic spindle rotates significantly (00:42-00:48) to realign the spindle along the internal apical – basal polarity axis and to ensure normal asymmetric cell division. “00:00” corresponds to telophase. Time scale is hh:mm and the scale bar is 3μm.

**File Name: Supplementary Movie 7**

**Description:** Neuroblast expressing GFP::Polo together with centriole tethered PACT::vhhGFP4.

Larval neuroblast expressing the transgene GFP::Polo (green), the microtubule marker mCherry::Jupiter (white) and the PACT::vhhGFP4 nanobody; both UAS lines are driven by the neuroblast-specific worGal4 transgene. Both MTOCs remain active and anchored to the apical cortex throughout interphase. Centrioles split only 6 minutes before mitosis starts (00:48). “00:00” corresponds to telophase. Time scale is hh:mm and the scale bar is 3μm.

## Methods

### Fly Strains, Transgenes and fluorescent markers

The following fly strains were used: Cnb RNAi (VDRC, 28651GD), *wdr62^⊗^ ^3-9^* allele ^19^, *Df(2L)Exel8005* (a deficiency removing the entire *wdr62* locus and adjacent genes; BDSC), *worniu*-*Gal4* ^39^, *pUbq-DSas-6::GFP* ^40^, *Cnn::GFP*, *Polo::GFP^CC01326^* (protein trap line) ^30^, *GFP::Polo* (genomic rescue construct using Polo’s endogenous enhancer) ^36^, *pUbq-Asl::GFP* ^41^, *pUbq-YFP::Cnb* ^11^, *YFP::Cnb^T4A,T9A,S82A^* ^11^, *nos-Cas9/Cyo* (BDSC), *y^1^, w^67c23^, P{y[+mDint2]=Crey}1b; D/TM3, Sb^1^* (BDSC), *y^1^, M{Act5C-Cas9.P.RFP-}ZH-2A, w^1118^, DNAlig4^169^* (BDSC), *worGal4, UAS-mCherry::Jupiter* ^16^, *cnb ^e00267^* ^18^, *Df(3L)ED4284* (*cnb* deficiency; BDSC), *polo^1^* ^42^, *polo^16-1^* ^43^, *pUASp-YFP::Cnb::PACT* ^18^.

The following mutant alleles and transgenes were generated for this paper: *Polo::EGFP*, *SspB::EGFP::Polo*, *Plp::EGFP*, *Cnb::EGFP*, *SspB::Dendra2::Cnb*, *mCherry::Cnb::PACT*, *PACT::HA::VhhGFP*, *SspB::mCherry*, *iLID::PACT::HA*, and *iLID::PACT::GFP*.

Unless specified otherwise, all strains were raised on standard medium at 25 °C, under a 12L:12D light cycle.

### Generation of transgenes using CRISPR/Cas9

*Plp::EGFP, Polo::EGFP, SspB::EGFP::polo, SspB::Dendra2::cnb, and Cnb::EGFP* were generated with CRISPR/Cas9 technology. Target specific sequences with high efficiency were chosen using the CRISPR Optimal Target Finder (http://tools.flycrispr.molbio.wisc.edu/targetFinder/), the DRSC CRISPR finder (http://www.flyrnai.org/crispr/), and the Efficiency Predictor (http://www.flyrnai.org/ evaluateCrispr/) web tools. Sense and antisense primers for these chosen sites were then cloned into pU6-BbsI-ChiRNA ^44^ between BbsI sites.

Plp::EGFP Target Site 1:

Sense:CTTCGAACTAGCGTCCACAAGGTC, Antisense:AAACGACCTTGTGGACGCTAGTTC

Plp::EGFP Target Site 2:

Sense:CTTCTGCTTATGGCTACATTTGGG, Antisense:AAACCCCAAATGTAGCCATAAGCA

Polo::EGFP Target Site 1:

Sense:CTTCGTCAGTCACCTCGGTGAATAT, Antisense AAACATATTCACCGAGGTGACTGAC

Polo::EGFP Target Site 1:

Sense:CTTCGAGACTGTAGGTGACGCATTC,Antisense:AAACGAATGCGTCACCTACAGTCTC

Cnb::EGFP Target Site 1:

Sense CTTCGCTCTATGAGACCTAAGCCT,Antisense AAACAGGCTTAGGTCTCATAGAGC

SspB::EGFP::polo Target Site 1:

Sense:CTTCGCTCTCCTTTCTTCTTTACTA, Antisense:AAACTAGTAAAGAAGAAGGAGAGC

SspB::Dendra2::cnb Target Site 1:

Sense:CTTCGGCAACCCTGTGCATCACCA), Antisense:AAACTGGTGAT GCACAGGGTTGCC)

To generate the replacement donor template SspB ^37^ (addgene #60416), the fluorophore (dendra2 or EGFP), and 1kb homology arms flanking the insertion site were cloned into pHD-DsRed-attP (Addgene plasmid # 51019) using Infusion technology (Takara/Clontech). Embryos expressing Act5C-Cas9 (BDSC#58492) for pHD-SspB::Dendra2::Cnb-DsRed, pHD-SspB::EGFP::polo-DsRed, or *nos-Cas9* ^45^ for polo::EGSP, plp::EGFP, and cnb::EGFP, were then injected with the replacement donor plasmid and its corresponding pU6-BbsI-ChiRNA. Injections were performed either in house or by Best Gene Injection Services (www.thebestgene.com). Successful events were detected by DsRed-positive screening in the F1 generation. Constitutively active Cre (BDSC#851) was then crossed in to remove the DsRed marker. Positive events were then balanced, genotyped, and sequenced.

### Generation of nanobody and optogenetic constructs

PACT::HA::vhhGFP4: The coding sequences of PACT ^33^ and vhhGFP4 ^34,35^ were PCR amplified and cloned into a pUAST-attB vector using In-Fusion technology (Takara, Clontech). The HA sequence was then added using overhang PCR. The resulting construct was injected into attP flies for targeted insertion on third chromosome (VK00027, BestGene).

mCherry::Cnb::PACT: The coding sequences of mCherry and Cnb::PACT were amplified by PCR (Cnb::PACT was amplified from pUASp-YFP::Cnb::PACT ^11^) and cloned into a pUASt-attB vector using In-Fusion technology (Takara, Clontech).The resulting construct was injected into attP flies for targeted insertion on the second chromosome (VK00018, BestGene).

SspB::mCherry: The coding sequence of SspB (addgene #60416) and mCherry were PCR amplified and cloned into a pUAST-attB vector using In-Fusion technology (Takara, Clontech). An AgeI site was added in the primers sequences to be inserted between SspB and mCherry. The resulting construct was injected into attP flies for targeted insertion on the third chromosome (VK00033, BestGene).

UAS-iLID::PACT::HA: The coding sequence of iLID (addgene #60411) and PACT ^33^ were PCR amplified and cloned into a pUAST-attB vector using In-Fusion technology (Takara, Clontech) along with a synthesized HA oligonucleotide sequence. The resulting construct was injected into attP flies for targeted insertion on the second chromosome (VK00018, BestGene).

UAS-iLID::PACT::GFP: The coding sequence of iLID (addgene #60411), PACT ^33^ and GFP were PCR amplified and cloned into a pUAST-attB vector using In-Fusion technology (Takara, Clontech). An XhoI site was added in the primers sequences to be inserted between iLID and PACT, and an AgeI site was added between PACT and GFP. The resulting construct was injected into attP flies for targeted insertion on the third chromosome (VK00020, BestGene).

### Immunohistochemistry

The following antibodies were used for this study: rat anti-α-Tub (Serotec; 1:1000), mouse anti-α-Tub (DM1A, Sigma; 1:2500), rabbit anti-Asl (1:500), rabbit anti-Plp (1:1000) (gifts from J. Raff). Secondary antibodies were from Molecular Probes and the Jackson Immuno laboratory.

96-120h (AEL; after egg laying) larval brains were dissected in Schneider’s medium (Sigma) and fixed for 20 min in 4% paraformaldehyde in PEM (100mM PIPES pH 6.9, 1mM EGTA and 1mM MgSO4). After fixing, the brains were washed with PBSBT (1X PBS, 0.1% Triton-X-100 and 1% BSA) and then blocked with 1X PBSBT for 1h. Primary antibody dilution was prepared in 1X PBSBT and brains were incubated for up to 2 days at 4 °C. Brains were washed with 1X PBSBT four times for 20 minutes each and then incubated with secondary antibodies diluted in 1X PBSBT at 4 °C, overnight. The next day, brains were washed with 1X PBST (1x PBS, 0.1% Triton-X-100) four times for 20 minutes each and kept in Vectashield H-1000 (Vector laboratories) mounting media at 4 °C.

### Super–Resolution 3D Structured Illumination Microscopy (3D-SIM)

3D-SIM was performed on fixed brain samples using a DeltaVision OMX-Blaze system (version 4; GE Healthcare), equipped with 405, 445, 488, 514, 568 and 642 nm solid-state lasers. Images were acquired using a Plan Apo N 60x, 1.42 NA oil immersion objective lens (Olympus) and 4 liquid-cooled sCMOs cameras (pco Edge, full frame 2560 × 2160; Photometrics). Exciting light was directed through a movable optical grating to generate a fine-striped interference pattern on the sample plane. The pattern was shifted laterally through five phases and three angular rotations of 60° for each z section. Optical z-sections were separated by 0.125 µm. The laser lines 405, 488, 568 and 642 nm were used for 3D-SIM acquisition. Exposure times were typically between 3 and 100 ms, and the power of each laser was adjusted to achieve optimal intensities of between 5,000 and 8,000 counts in a raw image of 15-bit dynamic range at the lowest laser power possible to minimize photobleaching. Multichannel imaging was achieved through sequential acquisition of wavelengths by separate cameras.

### 3D-SIM Image Reconstruction

Raw 3D-SIM images were processed and reconstructed using the DeltaVision OMX SoftWoRx software package (GE Healthcare; Gustafsson, M. G. L. 2000). The resulting size of the reconstructed images was of 512 × 512 pixels from an initial set of 256 × 256 raw images. The channels were aligned in the image plane and around the optical axis using predetermined shifts as measured using a target lens and the SoftWoRx alignment tool. The channels were then carefully aligned using alignment parameter from control measurements with 0.5 µm diameter multi-spectral fluorescent beads (Invitrogen, Thermo Fisher Scientific).

### Live cell imaging

72-120h (AEL; after egg laying) larval brains were dissected in Schneider’s medium (Sigma-Aldrich, S0146) supplemented with 10% BGS (HyClone) and transferred to 50 μL wells (Ibidi, μ-Slide Angiogenesis) for live cell imaging. Live samples were imaged on a Perkin Elmer spinning disk confocal system “Ultra View VoX” with a Yokogawa spinning disk unit and two Hamamatsu C9100-50 frame transfer EMCCD cameras. A 63x / 1.40 oil immersion objective mounted on a Leica DMI 6000B was used. Live cell imaging data shown in Figure 3, 6 and S7 was obtained with an Andor revolution spinning disc confocal system, consisting of a Yokogawa CSU-X1 spinning disk unit and two Andor iXon3 DU-897-BV EMCCD cameras. Either a 60x/1.4NA or 100X/1.4NA oil immersion objective mounted on a Nikon Eclipse Ti microscope was used.

### Fluorescence recovery after photobleaching (FRAP) experiments

The 488 nm laser line was targeted to regions of interests using Andor’s FRAPPA module. ROI’s measured ∼ 2μm × 2μm. Images were acquired every 30-60s after bleaching event. EGFP intensity before and after bleaching was measured using Imaris’ “Spot” tool.

### Optogenetics experiments

Crosses for optogenetics experiments were reared in the dark at 25°C. Offspring from these crosses were raised in the dark and dissected after 4 days using red filters to minimize ambient and blue light exposure. Optogenetic trapping or relocalization was performed using 10-20% of the 488nm diode laser (50mW) line.

### Centriolar Age Measurements

To determine centriolar age, Asl intensity was used as a reference. The contours of non-overlapping centrioles were drawn in ImageJ based on Asl signal and saved as XY coordinates. Using a custom-made MatLab code the total centriolar intensity, above background values determined by the expérimenter, for Asl were calculated in the drawn centriolar areas. Total Asl intensity was then used to determine centriolar age as daughter centrioles have lower intensity than mother centrioles. The same XY coordinates were used to measure total pixel intensity for markers of interest (e.g Polo::GFP, Plp::EGFP). Asymmetry ratios for markers of interest were then determined by dividing total daughter centriole intensity with total mother centriole intensity.

### Definition of statistical tests, sample number, sample collection, replicates

For each experiment, the data was collected from at least 2 independent experiments. All statistical details (replicates, n, statistical test used and p-values) for each experiment can be found in the corresponding figure legend. Statistical analyses were performed on Prism (GraphPad software). Statistical significance was assessed with a two-sided non-parametric Mann-Whitney test to compare ranks between two samples with variable variances and non-Gaussian distributions. P values < 0.05 were considered significant; *; p < 0.05 **; p < 0.01; ***; p < 0.001; ****; p < 0.0001.

### Computer codes

Custom made Matlab codes used for data analysis are available upon request.

### Data availability

The authors declare that the data supporting the findings of this study are available within the paper and its supplementary information files.

