## Supplementary figures and images for "Dynamic centriolar relocalization of Polo kinase and Centrobin in early mitosis primes centrosome asymmetry in fly neural stem cells"

### Supplemental Figure 1

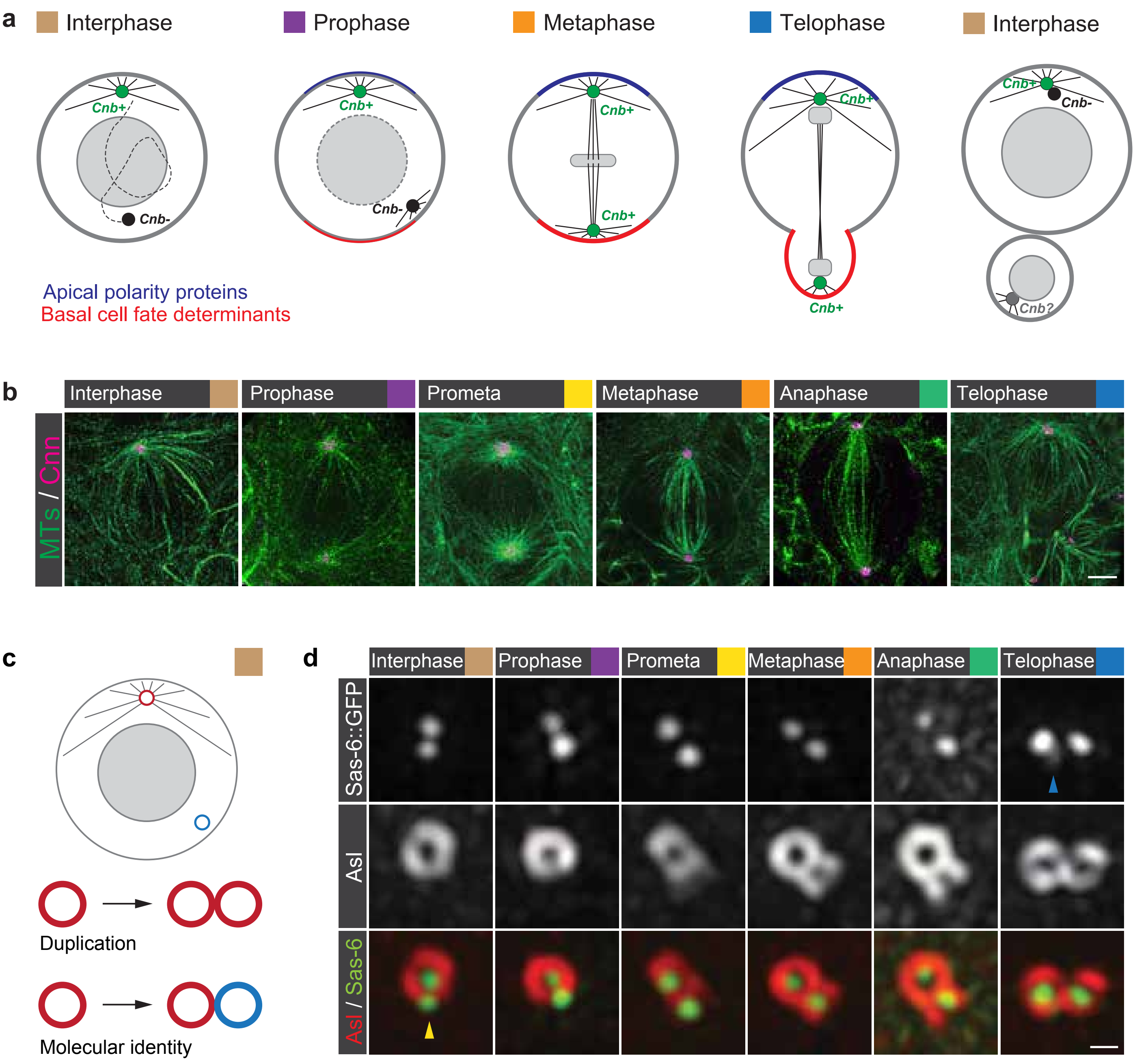

### Supplemental Figure 2

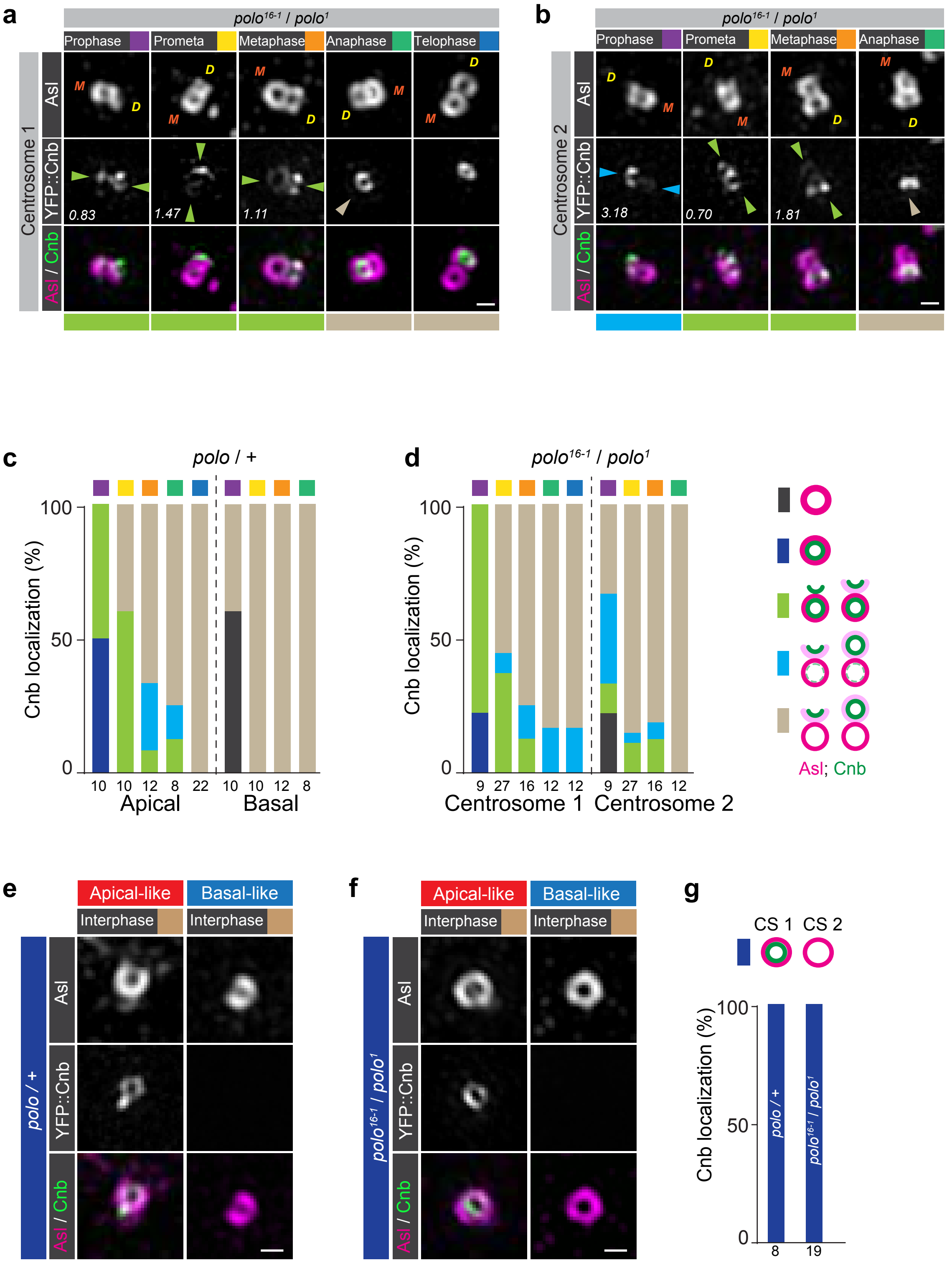

### Supplemental Figure 3

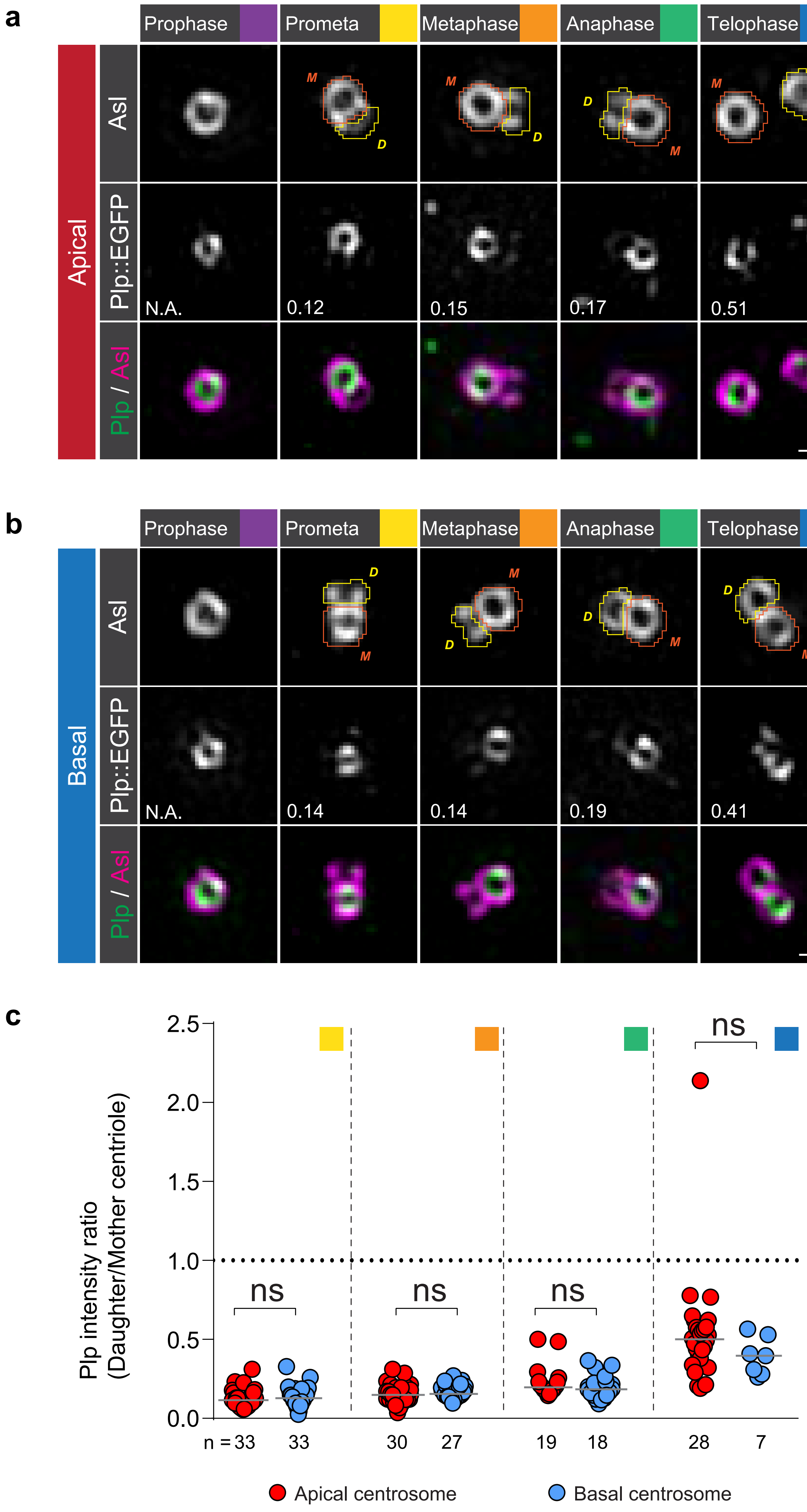

### Supplemental Figure 4

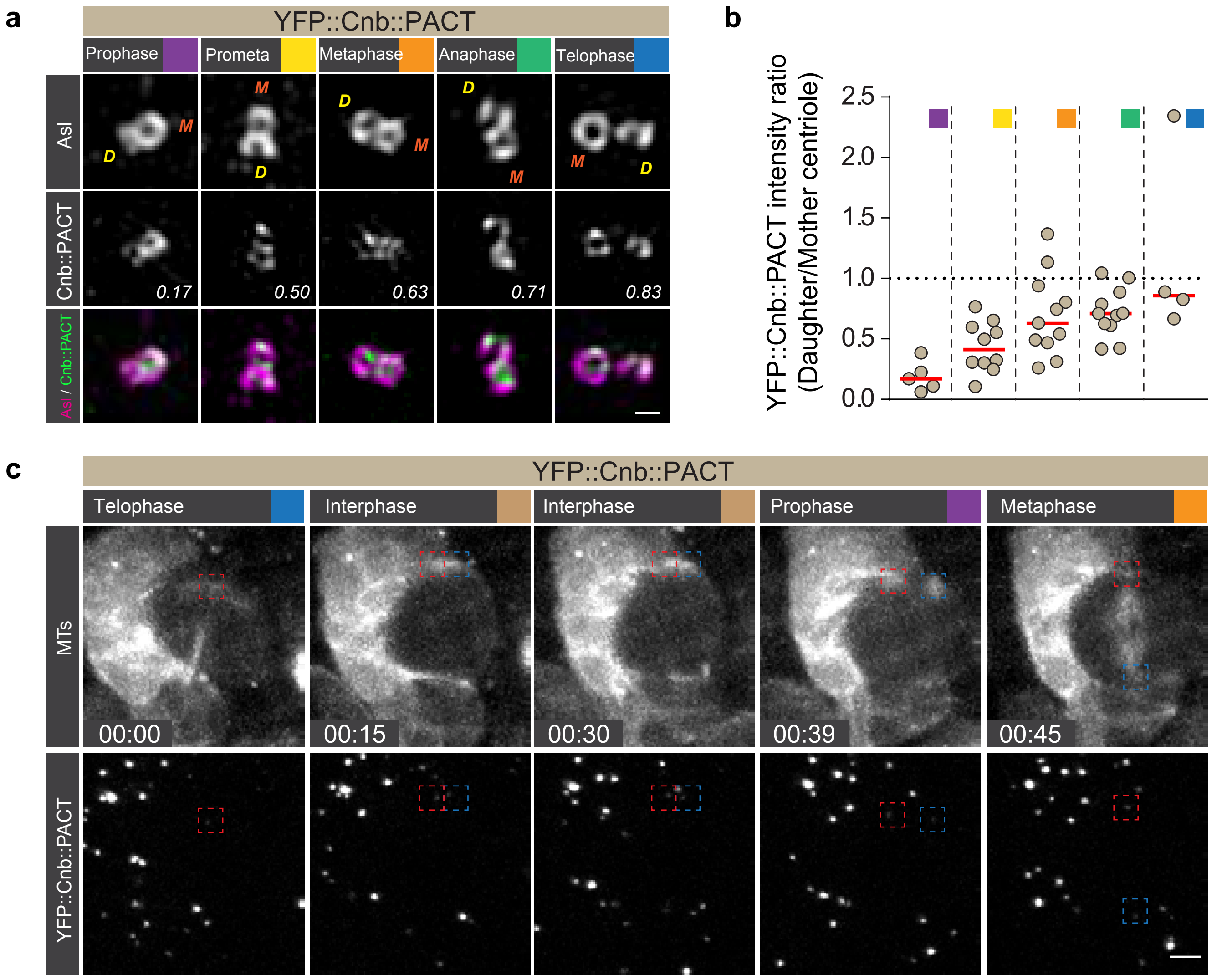

### Supplemental Figure 5

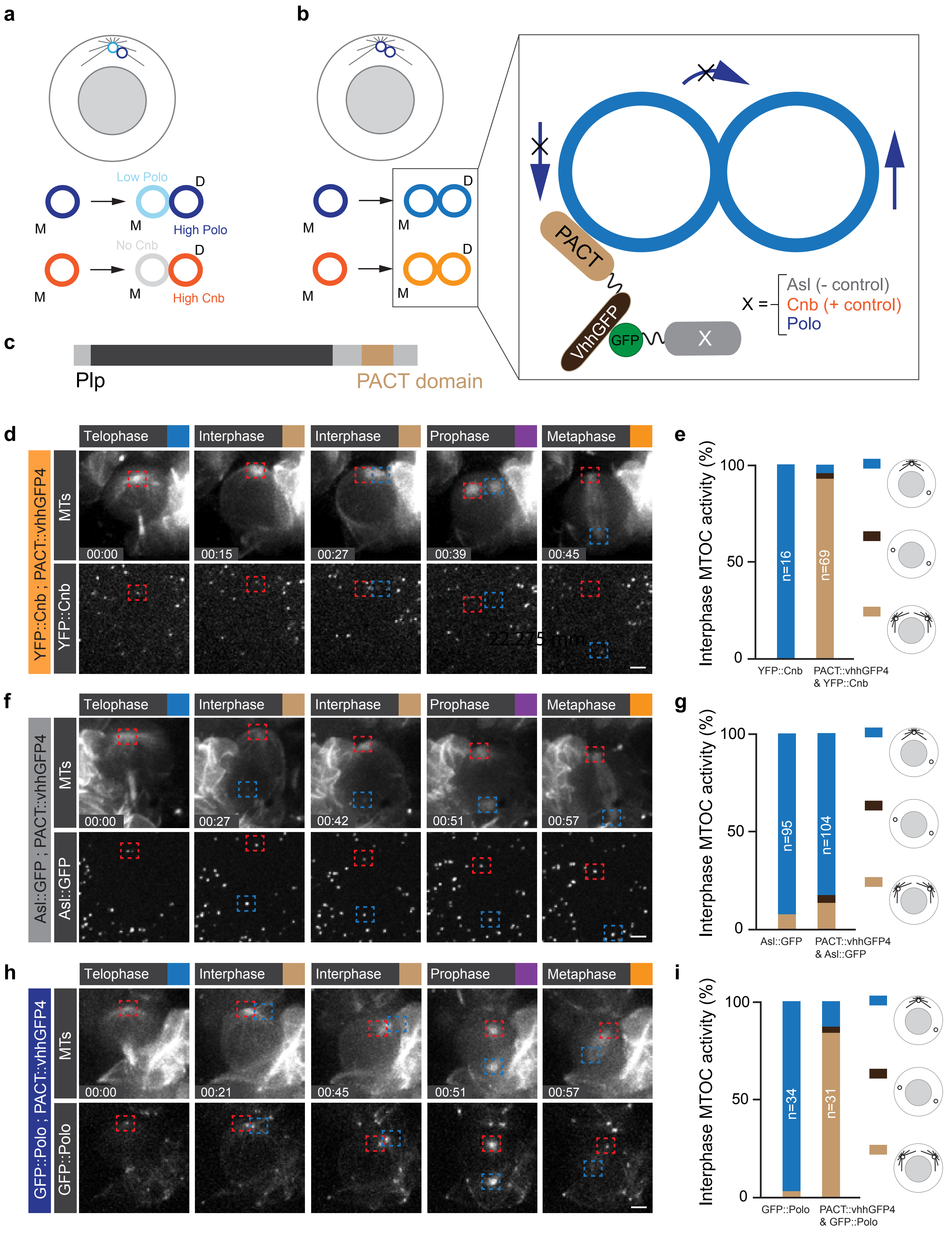

### Supplemental Figure 6

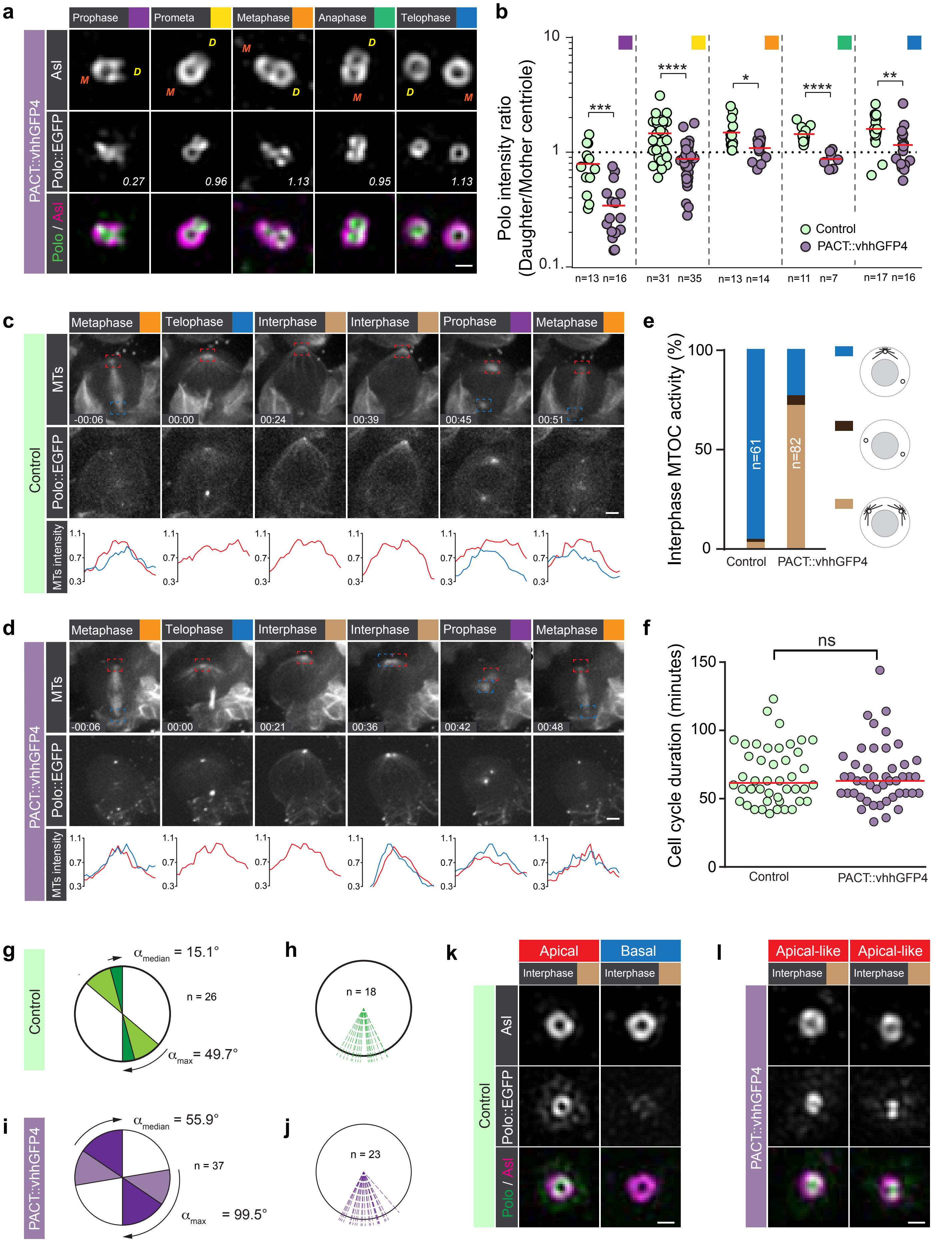

### Supplemental Figure 7

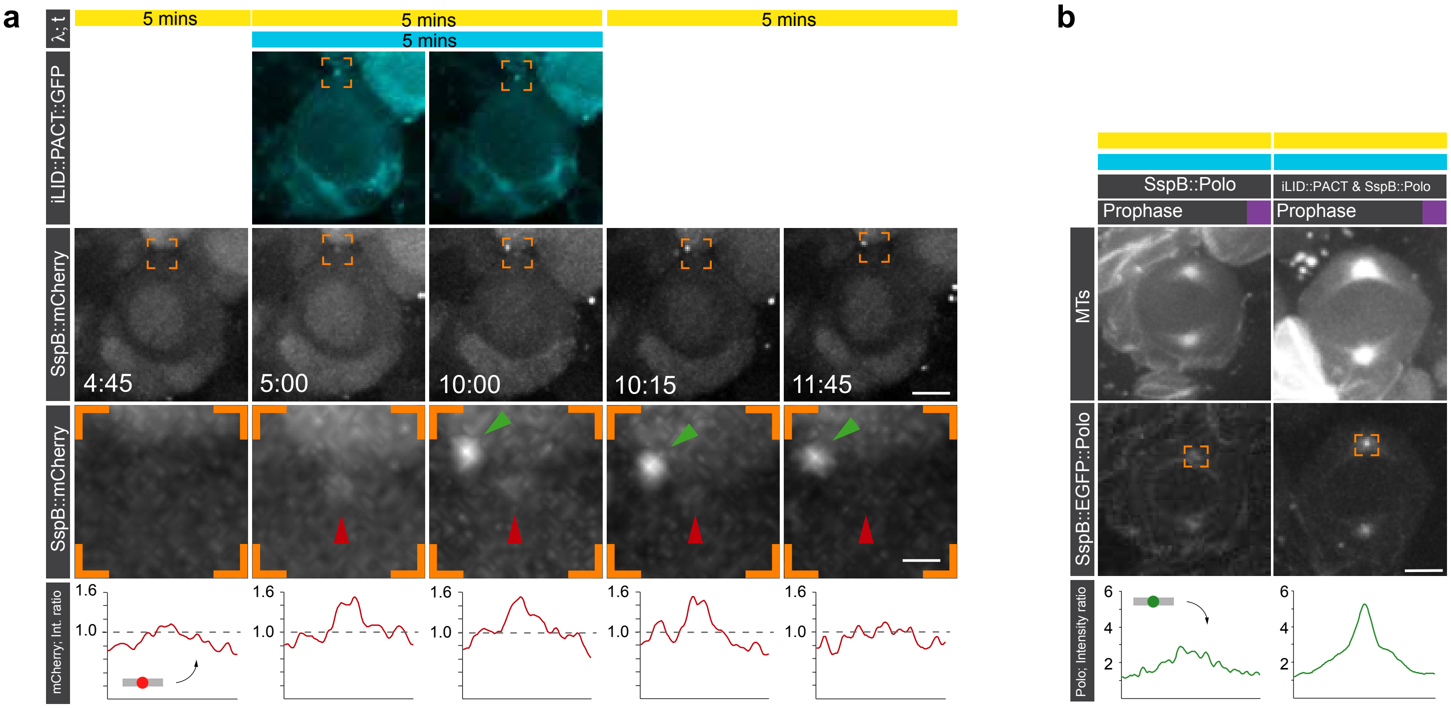
